# Conformational Tuning of Amylin by Charged Styrene-maleic-acid Copolymers

**DOI:** 10.1101/2020.04.23.057547

**Authors:** Bikash R. Sahoo, Christopher L. Souders, Takahiro W. Nakayama, Zhou Deng, Hunter Linton, Saba Suladze, Magdalena I. Ivanova, Bernd Reif, Toshio Ando, Christopher J. Martyniuk, Ayyalusamy Ramamoorthy

## Abstract

Human amylin forms structurally heterogeneous amyloids that have been linked to type-2 diabetes. Thus, understanding the molecular interactions governing amylin aggregation can provide mechanistic insights in its pathogenic formation. Here, we demonstrate that fibril formation of amylin is altered by synthetic amphipathic copolymer derivatives of the styrene-maleic-acid (SMAQA and SMAEA). High-speed AFM is used to follow the real-time aggregation of amylin by observing the rapid formation of *de novo* globular oligomers and arrestment of fibrillation by the positively-charged SMAQA. We also observed an accelerated fibril formation in the presence of the negatively-charged SMAEA. These findings were further validated by fluorescence, SOFAST-HMQC, DOSY and STD NMR experiments. Conformational analysis by CD and FT-IR revealed that the SMA copolymers modulate the conformation of amylin aggregates. While the species formed with SMAQA are α-helical, the ones formed with SMAEA are rich in β-sheet structure. The interacting interfaces between SMAEA or SMAQA and amylin are mapped by NMR and microseconds all-atom MD simulation. SMAEA displayed π-π interaction with Phe23, electrostatic π-cation interaction with His18 and hydrophobic packing with Ala13 and Val17; whereas SMAQA showed a selective interaction with amylin’s C-terminus (residues 31-37) that belongs to one of the two β-sheet regions (residues 14-19 and 31-36) involved in amylin fibrillation. Toxicity analysis showed both SMA copolymers to be non-toxic *in vitro* and the amylin species formed with the copolymers showed minimal deformity to zebrafish embryos. Together, this study demonstrates that chemical tools, such as copolymers, can be used to modulate amylin aggregation, alter the conformation of species.

## Introduction

The 37-residue hormone, islet-amyloid polypeptide (IAPP, or amylin) forms pathological amyloid inclusions in the pancreas of individuals with type-2 diabetes (T2D)[1], a devastating disease affecting millions of people. Normally, amylin is involved in the glycemic control and is co-secreted with insulin from pancreatic β-cells in response to food intake.[2] Amylin readily assembles to form amyloid fibrils causing β-cell dysfunction and death.[3] Evidence suggests that transient amylin species, precursors to amyloid fibril formation, are also associated with cellular toxicity, dysfunction and death. These transient species are typically small oligomers and have been shown to impair islet β-cell function and cause loss of β-cell mass.[1,4–7] Thus, modulating amylin aggregation with small-molecules to produce non-toxic species can be an useful strategy for controlling the disease progression.[8]

Many inhibitors have been developed by targeting different stages of fibril formation of amylin. For example, a common group of compounds, that inhibits amylin aggregation, stabilizes the water-soluble states of amylin monomers.[9–12] Small molecules, nanoparticles and polymers, have been shown to reduce the mass of cytotoxic oligomer species by rapid induction of non-crystalline fibers.[13–15] Studies have also exploited the strategy of targeting amylin monomers through electrostatic interactions to modulate its aggregation. For example, silver and iron nanoparticles were designed to sequester amylin monomers from fibers through electrostatic interactions.[16] Cationic and anionic polymers have been developed to either inhibit or accelerate amyloid aggregation.[9, 12] Similarly, surface-charge manipulation of quantum dots, gold or polymeric nanoparticles has been shown to effectively control amyloid aggregation.[17–20] Additionally, strong electrostatic interaction between amylin and anionic lipid membrane has been shown to yield the amylin species with distinct properties.[21, 22] Based on these reported studies, it is clear that fibril formation of amylin is strongly influenced by charge-charge interaction. Therefore, such interactions can be utilized in the development of a strategy to regulate amyloid formation by either inhibition or acceleration of fibril formation.[9,12,15,23]

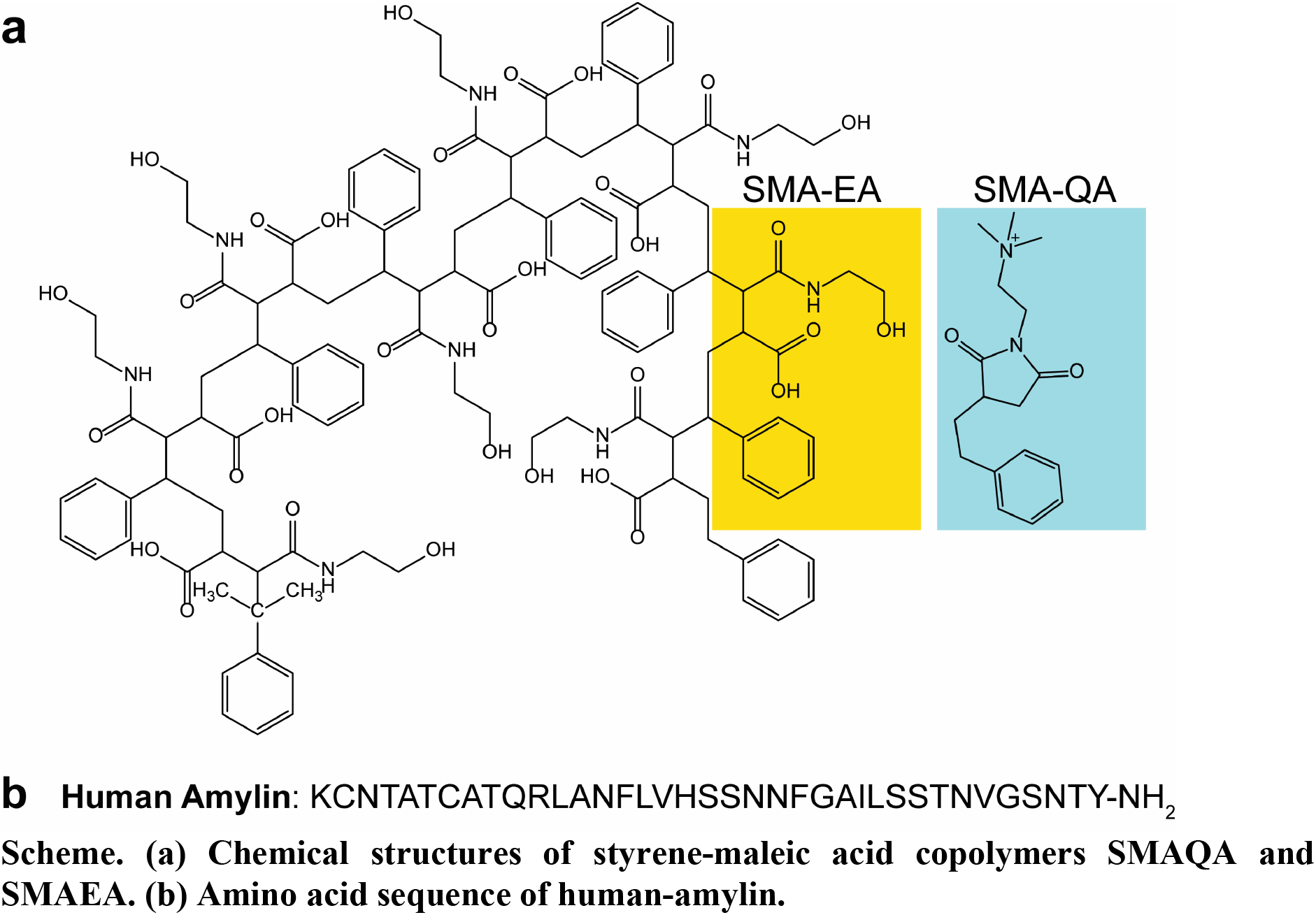

In this study, we utilize the charge-charge interactions with designed molecules[24, 25] to alter the course of amylin aggregation by tuning the conformation of the intermediate amylin species. For this purpose, we synthesized two different amphipathic styrene-based copolymers: SMAQA and SMAEA (Scheme **a**). These polymers are comprised of a conserved hydrophobic styrene moiety and a charged functional group that provide an overall positive charge for SMAQA and negative charge for SMAEA (Scheme **a**). Both SMAQA and SMAEA have been shown to form lipid-nanodiscs, and are used for the direct extraction of membrane proteins from cell-lysates.[24–27] Results reported in this study demonstrate that both SMAQA and SMAEA can be used to alter amylin fibril formation. Specifically, the presence of a positively-charged SMAQA rapidly converts monomers of amylin to globular oligomers, whereas the negatively-charged SMAEA accelerates fiber formation. Results obtained from cellular and zebrafish models show that the polymer-induced amylin species are mildly toxic and trend to be less toxic than amylin oligomers (fibrils and monomers) only.

## Results and Discussion

### Amylin fibril formation with SMAQA and SMAEA monitored by real-time HS-AFM

High-speed atomic force microscopy (HS-AFM) was used to monitor the real-time aggregation of amylin monomers by using sonicated amylin fibril as a seed (with seeds:monomer = 1:19 molar ratio) to accelerate the aggregation within the time-scale of HS-AFM observation. We observed both the *de novo* fiber formation and the growth of preformed fiber seeds (Figure 1, Video SV1). The fiber seeds grew fast on one end (indicated by the arrow-head) and slow on the opposite end (arrow-tail) after the addition of freshly prepared 5 *µ*M amylin monomers (Figure 1, Video SV1). The formation of *de novo* fibers was also observed at 1000, 1250 and 1350 s (marked with circles in Figure 1); this observation demonstrates the suitability of HS-AFM to capture *de novo* fiber formation, and the growth of *de novo* fibers and pre-seeded fibers in the sample at the same time. This type of seeded growth closely resembles the fibril formation of amyloid-β peptide as reported by a HS-AFM study.[28]

**Figure 1.**
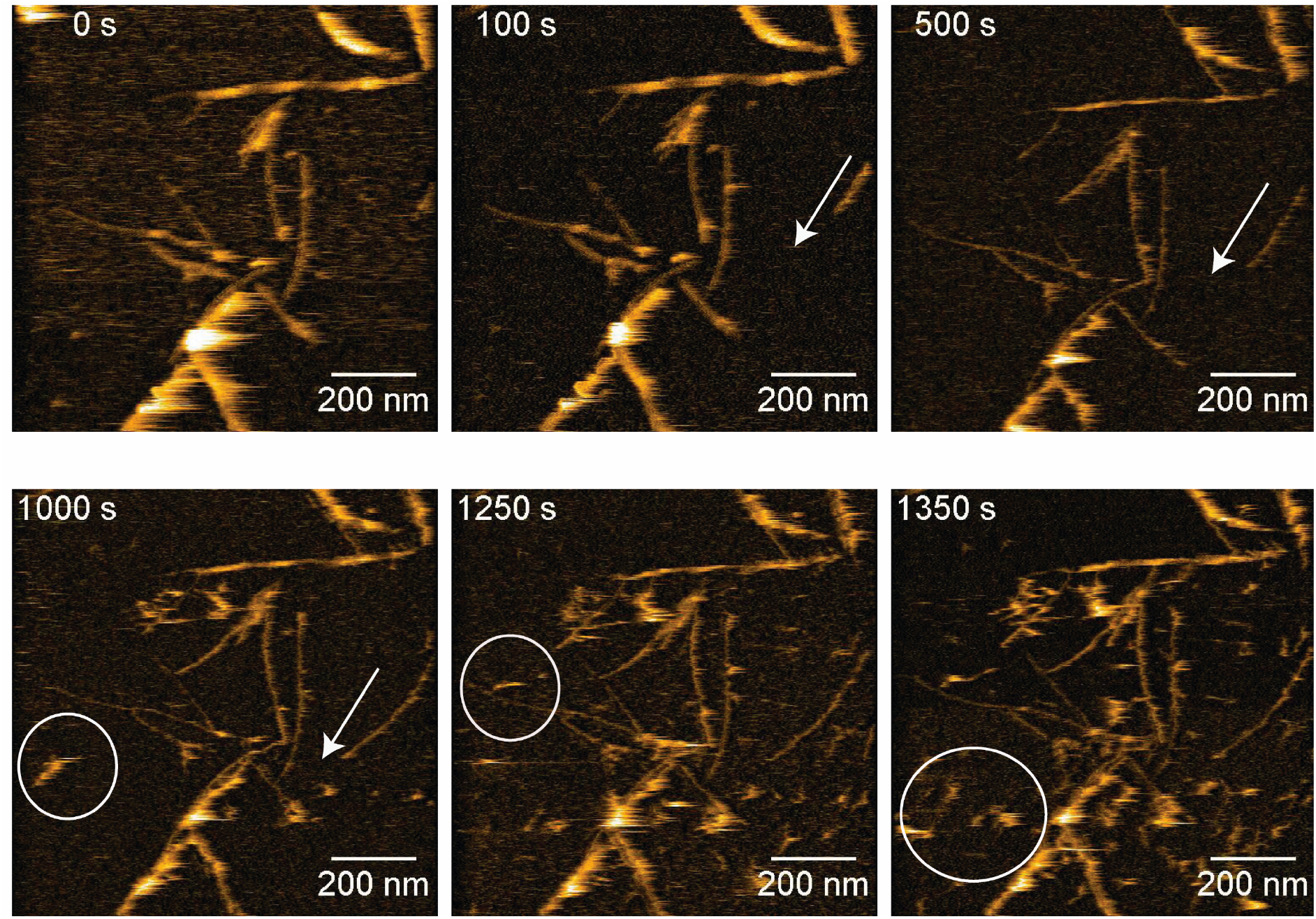
HS-AFM images showing the growth of human-amylin fibrils on a mica stage using freshly dissolved 5 *µ*M amylin monomers which was mixed with preformed fibrils deposited on the mica surface. Arrow indicates fast-growing (arrow head) and slow- growing (arrow tail) ends of a selected amylin fiber seed. The *de novo* formed fibers are circled.

The effect of each of the polymers on amylin fibrilation was next examined in real-time by HS-AFM. As shown in Figure 2, *de novo* globular amylin species (≈18 nm in diameter) was observed after the addition of 11 *µ*g/mL SMAQA (Figure 2 (a and c), see the Video SV2 at 225 s). While the number of globulomers increased with time as shown in Figure 2(a and e) and in the Video SV3 (from 500 to 1335 s), we observed that that the initially added fibril-seeds did not grow (Figure 2a). In addition, the globulomers were found to grow in size but did not convert into fibers within the measurement time scale (∼1,350 s), as shown in Figure 2e and video SV2. In contrast to SMAQA, in the presence of 11 *µ*g/mL SMAEA, the initially added amylin fibril-seeds grew in size and the *de novo formed* globulomers were converted into fibrils (Figure 2(g,i) and Video SV6). In addition, the fibril growth of amylin was bidirectional which is similar to the fibril growth in the absence of copolymers (Figure 1, Videos SV4 and SV5). Kymographs of fibrillation of the selected fiber in the presence of SMAEA showed that the ends of fibrils grew 40 to 50 nm within a time-interval of 200 s (Figure 2h, see Video SV4).

**Figure 2.**
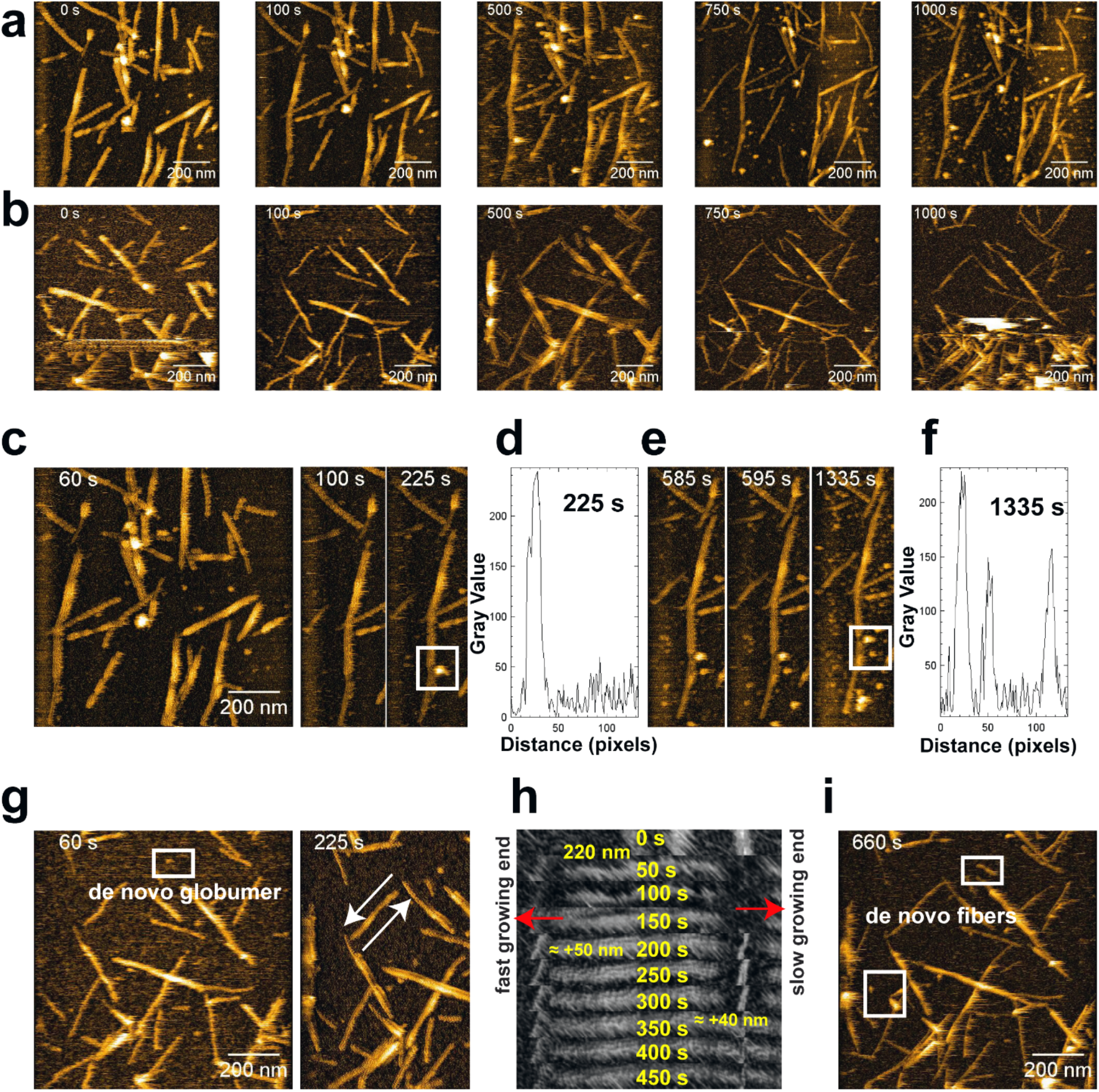
The effect of SMA copolymers on amylin fibrillation monitored by real-time HS-AFM. HS-AFM images of 5 μM amylin dissolved in 30 mM NaAc, pH 5.5 in the presence of 11 *µ*g/mL SMAQA (a) and SMAEA (b). The *de novo* globulomer formation in the presence of SMAQA is shown inside a white box (in (c) and (e)). The size of the globulomers (as shown in (d) and (f)) were determined using ImageJ where on the x-axis 1pixel=2 nm. The bidirectional fibril growth in the presence of SMAEA is shown by arrows at t=225 s (g), and the time lapse images of fast and slow growing ends. (h) The *de novo* globulomer formation is shown inside a white box in (g). The growth of *de novo* globulomer into fibers in the presence of SMAEA is shown inside the white box in (i).

### Opposite effects of SMAQA and SMAEA copolymers on amylin fibrillation

In order to investigate the effect of the polymer on the kinetics of amylin aggregation, thioflavin-T (ThT) based fluorescence experiments were carried out. We first determined that 11 *µ*g/mL of SMAQA co-polymer alone does not interfere with ThT fluorescence (Figure S1), whereas an increased ThT fluorescence was observed for 11 *µ*g/mL SMAEA co-polymer alone. Then, ThT fluorescence of matured amylin fibers alone and with SMAEA was measured to evaluate the contribution from SMAEA alone (Figure 3a). As shown in Figures 3(a,d) and S1, freshly dissolved amylin monomers, or the matured amylin fibers, mixed with SMAEA presented a substantial increase in ThT fluorescence as compared to SMAEA alone. We further examined the ThT fluorescence of samples that contained 1% to 7% (v/v) mature fibers (prepared from 100 *µ*M amylin monomers) and 11 or 22 *µ*g/mL copolymer (Figure 3a). Mature amylin fibrils were added at the following time intervals: t=0 to 10 minutes – no fibrils, t=10-20 minutes – 1% fibrils, t= 20-30 minutes – 2% fibrils, and t= 40-100 minutes – 7%. Results from these samples showed that SMAEA contribution to ThT fluorescence is much smaller when compared to that from amylin fibers (Figure 3a-c). Therefore, we concluded that ThT can be used as a molecular probe to monitor amylin fibrillation in the presence of either of the co-polymers in real-time.

**Figure 3.**
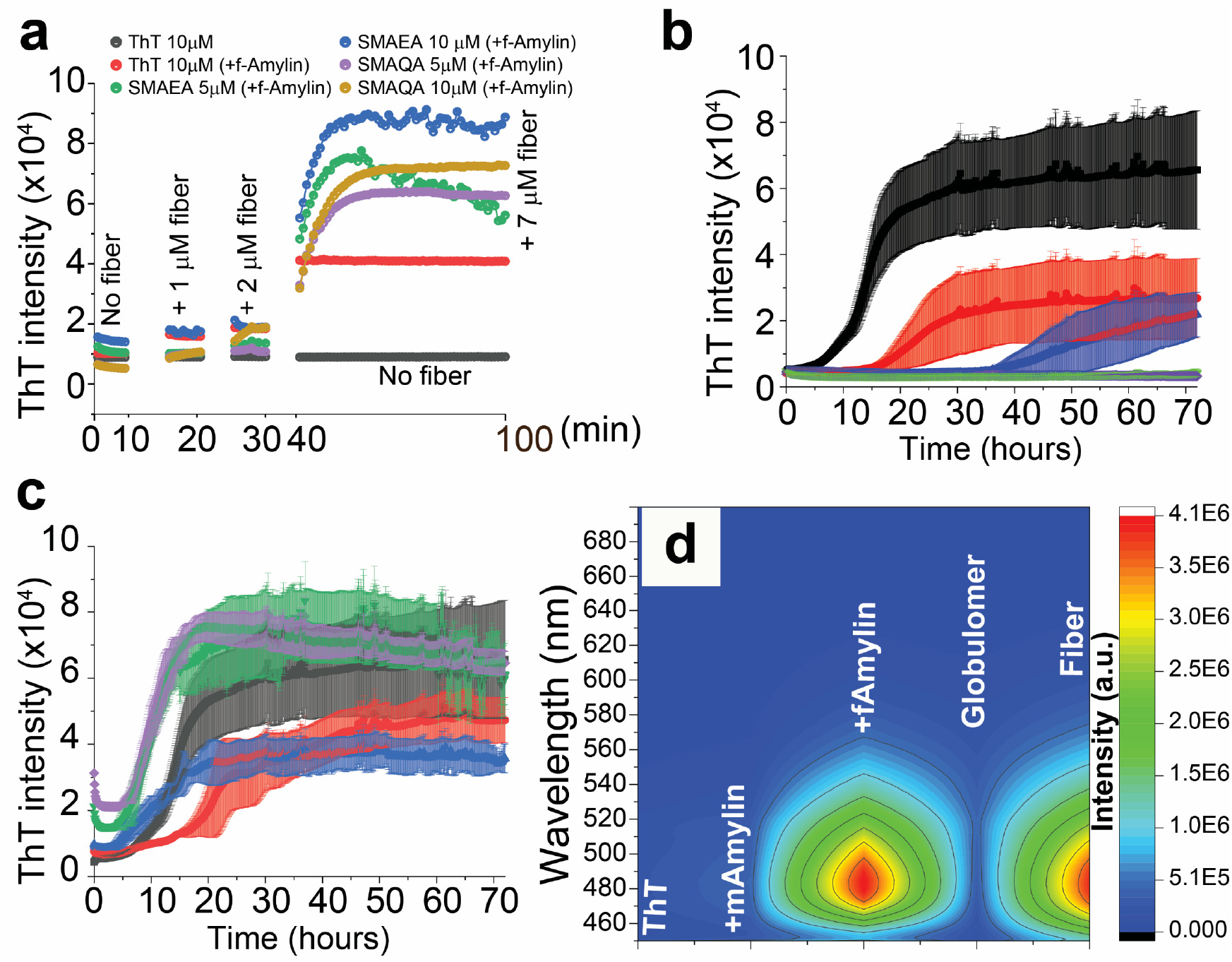
(a) Monitoring the change in thioflavin-T (10 µM) fluorescence intensity in the presence of SMA copolymers as a function of amylin fiber concentration (see methods). Fluorescence intensity of samples containing no amylin fiber is measured in the initial 10 minutes. Preformed amylin fibers (1%, 2% and 7% (v/v)) (except the black curve for ThT alone) were added at time-intervals 10, 20 and 40 minutes, respectively. (b-c) Thioflavin-T fluorescence spectra of 10 µM human-amylin (4 replicates) dissolved in NaAc buffer, pH 5.5 in the absence (black) or presence (red: 5.5 µg/mL; blue: 11 µg/mL; green: 22 µg/mL; purple: 44 µg/mL) SMAQA (b) or SMAEA (c). (d) 2D plot showing the fluorescence of 10 µM of ThT (only), and ThT mixed with 5 µM amylin monomers (labeled as mAmylin), preformed fibers (fAmylin), filtered SMAQA (11 µg/mL)+ 5 µM amylin (labeled as globulomer), and filtered SMAEA (11 µg/mL)+ 5 µM amylin (labeled as Fiber) (see Figure S1).

The effect of different concentrations of SMA copolymers on amylin aggregation was next examined. Increase in ThT fluorescence was observed when amylin monomers were incubated with SMAEA indicating the formation of amyloid fibers (Figure S1). On the other hand, no change in ThT fluorescence was observed when amylin was co-incubated with SMAQA suggesting the formation of non-fibrillar species (Figures S1 and 3(b,c)). While a low concentration (5.5 µg/mL) of SMAEA or SMAQA delayed the aggregation of amylin (monomer concentration of 10 µM), acceleration by SMAEA (Figure 3c) and increasing delay (or inhibition) by SMAQA (Figure 3b) were observed at higher concentrations.

Since we observed a quick fibrillation of amylin (within few minutes) in the presence of SMAEA and globulomers in the presence of SMAQA (within few minutes) by HS-AFM experiments (Figures 1 and 2), static fluorescence experiments were performed to monitor ThT binding within few minutes after the addition of the polymer to amylin monomers. These experiments showed no increase in the fluorescence intensity for a sample of amylin monomers mixed with SMAQA. On other hand, a strong fluorescence was observed for a sample containing SMAEA and amylin monomers (Figure 3d), which is similar to the ThT binding with matured fibers.

Fluorescence experiments were also performed by adding ThT dye to the large amylin aggregates that were formed in the presence of polymer and obtained by filtration as explained in the Methods section. A very weak fluorescence for SMAQA induced amylin globulomers and a strong fluorescence for SMAEA induced amylin fibers were observed (Figures 3d and S1). These ThT based fluorescence results are in agreement with HS-AFM results and suggest the fibrillar and non-fibrillar amylin aggregates formed in the presence of SMAEA and SMAQA, respectively.

### Rapid and distinct conformational alteration in amylin by SMAEA and SMAQA

To better understand the above described differences on the effects of polymers on amylin aggregation, CD experiments were performed to probe amylin’s secondary structural changes in the presence of each of the polymers. Freshly dissolved amylin in NaAc buffer, pH 5.5 showed a significant change in the secondary structure in the presence of 11 or 27.5 *µ*g/mL SMAEA (which is below the critical micellar concentration of the polymer) and no change in the presence of SMAQA (Figures 4a, S2 and S3). The analysis of CD spectra of 25 *µ*M amylin (day-1, ∼15 minutes) in the absence of SMA copolymers using BestSel [29] presented 17.2, 8.6 and 5.4 % of α, antiparallel β and parallel β structures (experimental vs fitted NRMSD (normalized root mean square deviation) ∼0.01). On the other hand, 25 *µ*M amylin mixed with 55 *µ*g/mL SMAEA (day-1) shifted the CD minimum from ≈200 to ≈227 nm (Figure 4a, blue) indicating a predominant β-sheet rich structure characterized with ∼4.6% α-helix and ∼27.8% antiparallel β-sheet structures (experimental vs fitted NRMSD ∼0.03). The presence of 55 *µ*g/mL SMAQA amylin showed an α-helix conformation (Figure 4a, green) within ∼15 minutes of incubation characterized with 36.3% α-helix and 13.4% antiparallel β-sheet (experimental vs fitted NRMSD ∼0.03). Similar to amylin-only samples, amylin samples mixed with SMAEA that were incubated for ∼24-hour showed a decrease in molar ellipticity indicating amylin fibrillation (Figure 4a).[12] In contrast, SMAQA-amylin mixture presented a α/β mixed structure characterized with ∼32.9% α-helix, ∼14.5% antiparallel and ∼11.5% parallel β-sheets (based on the analysis using BestSel[29]) with a little change in molar ellipticity upon ∼24-hour incubation (Figure 4a). Thus, the CD results suggest that the distinct structural changes observed for amylin in the presence of SMAEA or SMAQA are likely due to electrostatic interactions of amylin with polymers. It should be noted that the disordered to helical structural transition observed in the presence of SMAQA in this study was also observed for amylin in the presence of a positively charged quaternary ammonium containing small molecule inhibitors and also in dodecylphosphocholine micelles, which suggest the common role of electrostatic interaction of amylin with positively charged molecules.[30, 31] Similarly, the anionic SMAEA that accelerates amylin fibrillation correlates to the effect of a negatively charged star-polymer that was reported to promote amylin aggregation.[9] It should also be noted that the polymers themselves do not show a characteristic CD spectrum in the absence of amylin (Figure S4), but they may interfere with CD signals from amylin when bound to amylin. Therefore, the CD observations were further verified using FT-IR experiments (Figure 4b).

**Figure 4.**
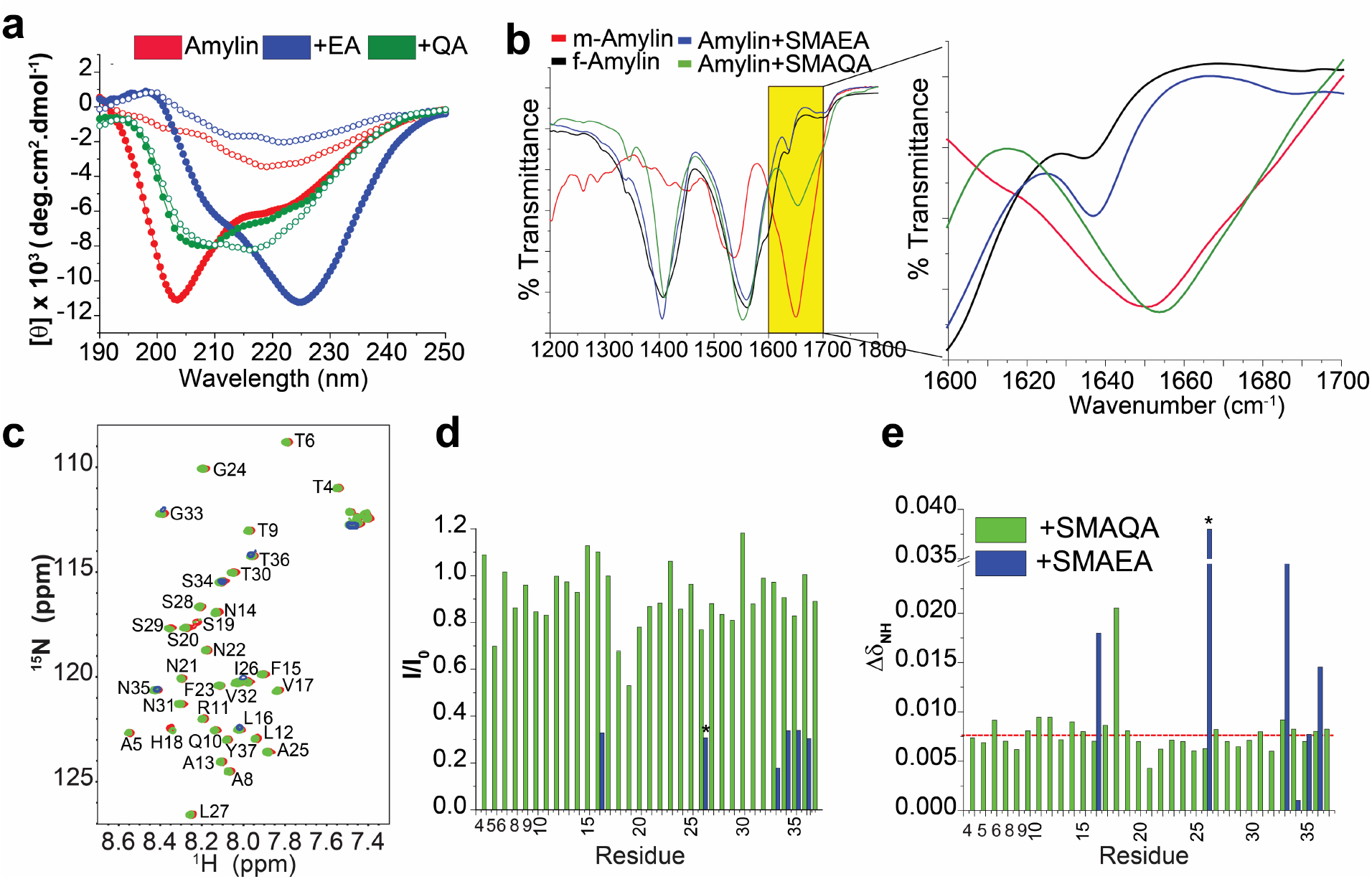
(a) Time-lapse far-UV CD measurement of 25 µM amylin dissolved in NaAc buffer, pH 5.5 in the presence and absence of 55 µg/mL SMAEA or SMAQA (∼15 min:solid circle; ∼24 h:open circle) (b) FT-IR spectra show the effect of copolymers on the secondary structure of human-amylin. 25 µM of freshly dissolved human-amylin mixed with 55 µg/mL SMAQA or SMAEA was incubated for ∼12-hours following lyophilization and used for FT-IR measurement. Region spanning 1600-1700 cm^−1^ representing the peptide secondary structure is highlighted and zoomed (box). Freshly dissolved amylin monomers and fibers prepared from 25 µM monomers following ∼72 hours continuous agitation at room temperature were lyophilized and used as a control for comparative conformational analysis. (c) 2D ^15^N/^1^H SOFAST-HMQC spectra of 25 µM ^15^N-labeled amylin in the absence (red) and presence of 55 µg/mL SMAQA (green) or SMAEA (blue) recorded on an 800 MHz NMR spectrometer at 25 °C. (d) Peak intensities measured from SOFAST-HMQC spectra: peak intensities of amylin residues in the absence (I_0_) and in the presence of polymer (I): SMAEA (blue) and SMAQA (green). * indicates peak intensities from I26/V32 in the SMAEA-amylin sample. (e) Chemical shift perturbations (CSPs) calculated using 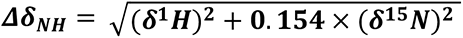. The dashed red line represents the average (CSP_avg_) value calculated for the amylin-SMAQA mixture, whereas CSP_avg_ is not shown for amylin-SMAEA due to significant loss of signal intensity for most of the residues.

FT-IR spectra of amylin monomers in the absence and presence of copolymers, and that of matured amylin fibers, are shown in Figure 4b. The amide-I absorption band revealed the backbone conformation of amylin without interference from the copolymer: a β-sheet conformation for the SMAEA-amylin complex (∼1637 cm^-1^, spectrum resembles that of the matured amylin fibers) and a partially folded helical conformation for the SMAQA-amylin complex (∼1653 cm^-1^, spectrum resembles that of amylin monomers ∼1649 cm^-1^) (Figure 4b). Taken together, the CD and FT-IR results confirm a distinct conformational state of amylin in the presence of the polymers when compared with freshly prepared amylin monomers. Further characterization of the amylin species by transmission electron microscopy (TEM) showed a spherical morphology in the presence of SMAQA and fibrillary morphology in the presence of SMAEA (Figure S5). We note that such spherical amylin species characterized by predominant α-helix conformation has also been observed previously.[32, 33]

### SMAEA drives amylin fibrillation via rapid β structure induction

We previously demonstrated a rapid β-sheet structure induction in Aβ1-40 upon its interaction with an oppositely charged polymer.[12] Therefore, we hypothesized that the charge-charge interaction between the cationic amylin (amylin: ∼+3 at pH 5.5) and anionic SMAEA could drive the rapid β-sheet structure induction observed in the CD results presented above. To test this hypothesis, the effect of SMAEA (55 µg/mL) on amylin (25 µM) structure was measured in a buffer solution at pH 8.5 (where amylin’s net charge is ≈0) or in a buffer containing 0.25 M NaCl. Interestingly, CD results showed that the change in buffer pH (from 5.5 to 8.5), or the addition of 0.25 M salt, interfered with the SMAEA-induced rapid β-sheet formation and yielded a α-helical structure for amylin (Figure S6 (a,b)). The observed CD spectrum of the amylin+SMAEA mixture in basic condition (or in the presence of salt) closely resembles with that of the SMAQA+amylin mixture which showed a α-helical conformation for amylin (Figure 4a, green). In addition, fluorescence assay showed that unlike SMAEA-induced amylin aggregation in the absence of salt (Figure 3b, green), a varying buffer ionic strength (10 mM to 1M NaCl) delayed amylin (10 µM) aggregation in the presence of 22 µg/mL SMAEA (Figure S6(c,d)). Whereas a varying concentration (10 mM, 50 mM, 100 mM and 1 M) of NaCl showed a decrease in the lag-time of amylin aggregation (Figure S6c). These results indicate that SMAEA is able to delay amylin’s aggregation even when the salt concentration is below the physiological condition (i.e. 150 mM). These results also suggest that the amylin aggregation mechanism and the interaction between amylin and SMAEA under physiological condition could be a complex process that depends on the concentration of salt, SMAEA and amylin.

### Molecular interaction between copolymers and amylin

To obtain atomic-resolution insights into the molecular mechanism underlying the interaction of SMA copolymers with amylin, NMR experiments and molecular dynamics (MD) simulation were carried out. 2D (^1^H/^15^N) SOFAST-HMQC spectrum shows dispersed resonances for 25 µM amylin (Figure 4c, red); the resonance assignment is based on a previously reported study.[34] Titration of 55 µg/mL SMAQA resulted in a decrease in the intensity for most of the resonances observed in the NMR spectrum (Figure 4(c and d), green). Since the measurement time for the 2D NMR spectrum is ≈ 1 hour, the loss in the signal intensity indicates an increase in the size of amylin species in the presence of SMAQA in agreement with the results from HS-AFM (Figure 2c), fluorescence (Figure 3) and TEM (Figures S5b). In particular, ≈>25% decrease in the signal intensity was observed for residues A5, H18, S19 and S20. Chemical shift perturbation (Δδ_NH_) analysis identified that several N- and C-terminal residues exhibited Δδ_NH_ above the average value (Figure 4e). On the other hand, when amylin was mixed with 55 µg/mL SMAEA, most of the resonances in the 2D spectrum disappeared except L16 and I26/V32. Residues spanning 33-36 exhibited a significant reduction in signal intensity (>75%) and chemical shift perturbation (Figure 4c-e, blue). The loss of NMR signals correlates well with amylin fibrillation observed from HS-AFM and fluorescence experiments (Figures 2e and 3c); amylin fibrils are usually not detectable in solution NMR experiments due to their immobile (or very slowly tumbling) nature in the time scale of NMR spectroscopy. The few peaks observed in the NMR spectrum likely to be from the flexible regions of the SMAEA-induced amylin species.

The time-lapsed ^1^H NMR experiments further revealed a significant loss of amylin (25 µM) amide-NH resonances (spanning ≈7.5 to 8.5 ppm) immediately after titration with SMAEA (55 µg/mL) (Figure 5(a,b)); note that the NMR measurement time-scale was <30 minutes. Visual inspection showed a turbid and precipitated SMAEA-amylin sample on day-7, whereas the SMAQA-amylin sample was found to be transparent (Figure 5, top center). Notably, the addition of either of the SMA copolymers rapidly induced amylin oligomers as indicated by the peak near ≈ -0.5 ppm (marked with * in the spectrum) which disappeared only in the sample containing SMAEA on day-7 as the oligomers would have been converted to fibers (Figure 5(a,b)). ^1^H NMR measurements of polymer-induced amylin globulomers (25 µM amylin and 55 µg/mL SMAQA) isolated using a 30-kDa filter (see methods) showed the absence of SMAQA (Figure S7). This was further validated using saturation-transfer-difference (STD) NMR measurements on the amylin+SMA mixed samples as shown in Figure 5(c,d). Saturation of amylin’s oligomer peak (≈-0.5 ppm, blue spectra) do not show a transfer of magnetization from amylin species to SMAEA/SMAQA polymer indicating that the polymers disassociate upon amylin aggregation. Saturation of -NR_3_^+^ protons (3.078 ppm, pink) of SMAQA showed a weak magnetization transfer to aromatic styrene protons resonating at ≈7 ppm (Figure 5c, pink), but no magnetization transfer to amylin was observed. Further, saturation of protons in the styrene-amide overlapping region (≈7.2 ppm) showed no detectable magnetization transfer. These STD NMR results are supported by the diffusion ordered spectroscopy (DOSY) measurements that showed a heterogeneous mixture in the amylin+SMA solution composed of large and small particles. A diffusion constant of ∼2×10^-11^ m/s^2^ was calculated for the SMAQA-induced amylin oligomers that are smaller (hydrodynamic diameter ≈30 nm) as compared to previously reported amylin oligomers in the absence of any additives (Figure S8a).[35] In addition, a self-diffusion constant of ∼1×10^-10^ m/s^2^ was calculated for SMAQA (-NR^3+^) suggesting the presence of free SMAQA in the solution. Similarly, amylin mixed with SMAEA solution also showed a diffusion constant value of ∼1×10^- 10^ m/s^2^ (Figure S8b) indicating the presence of fast diffusing polymers that do not associate with amylin aggregates.

**Figure 5.**
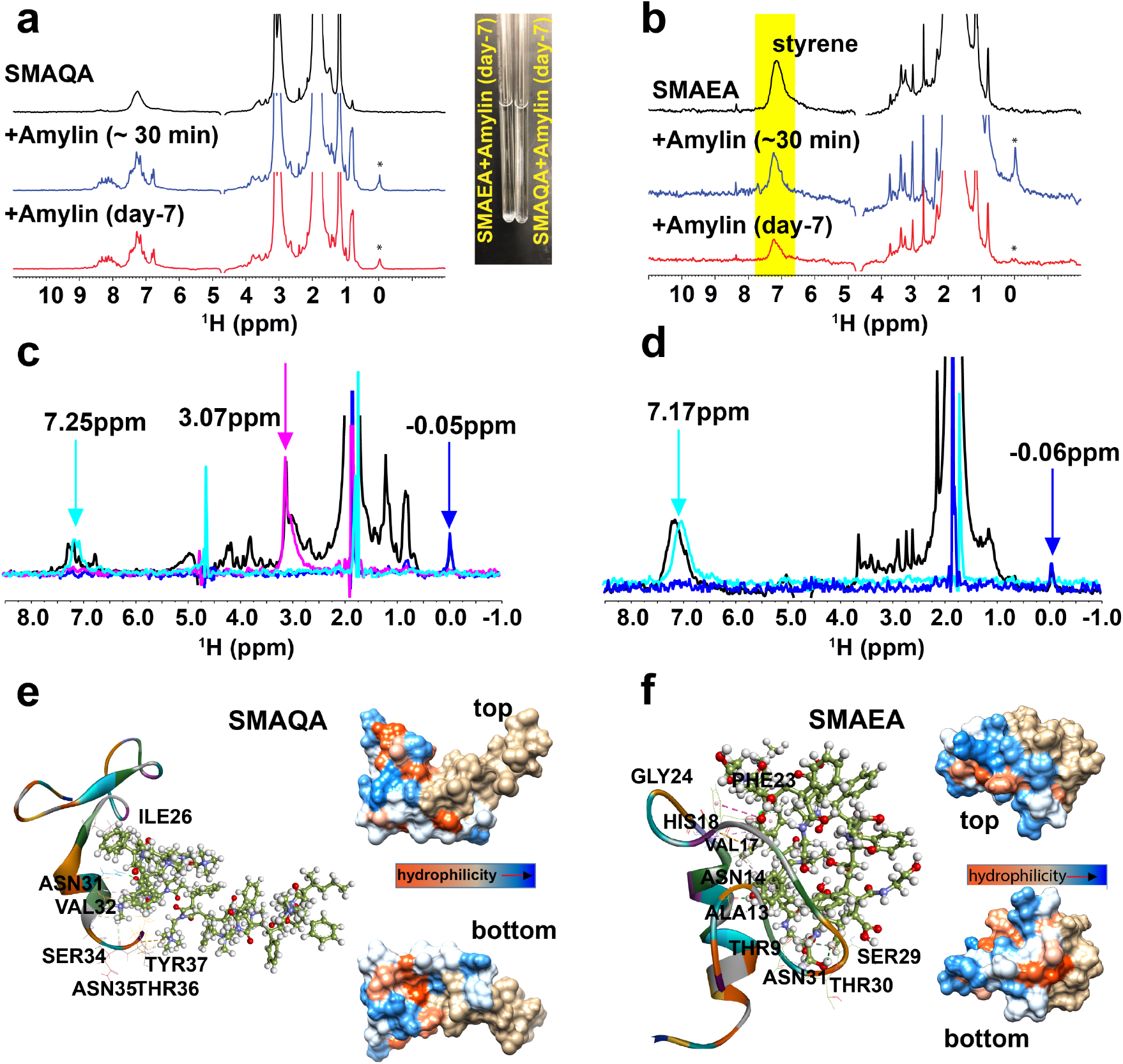
Structural interactions between amylin and SMA copolymers. (a-b) Time-lapse ^1^H NMR spectra of 55 µg/mL polymer (black) mixed with 25 µM amylin (blue and red) recorded on a 500 MHz NMR spectrometer at 25 °C at the indicated times. The styrene ^1^H peak is highlighted in yellow in (b). A broad low intensity peak from amylin oligomers appeared near ≈-0.5 ppm is indicated with *. The SMAEA-amylin solution showed precipitation whereas the SMAQA-amylin samples were clear at the end of day-7 as shown in the NMR tubes (top, center). (c-d) STD ^1^H NMR spectra of 25 μM amylin in the presence of 55 µg/mL SMAQA (c) and SMAEA (d) recorded on a 500 MHz NMR spectrometer at 25 °C. The reference spectrum is shown in black and the saturation transfer difference spectrum is shown in the indicted colors (blue or pink) and the saturated peaks are indicated by arrows. x-axis offset is used for the cyan spectrum to avoid spectral overlapping and for visual clarity. Snapshots obtained at 500 ns MD simulations are shown for SMAQA-amylin (e) and SMAEA-amylin (f) complexes. The interaction between polymer (ball-stick) with amylin (cartoon) shown in the structures that were generated using DSV v17.2.0. The amylin binding residues that form hydrogen bonding, electrostatic, hydrophobic and π-π interactions are labeled (shown as dashed lines). The top and bottom views of the hydrophobic surface maps of the complexes are shown on the right of the respective complex structure.

### Structure and dynamics of the SMA-amylin complex

The interaction between SMA copolymers and amylin was studied using all-atom MD simulations. Simulation results presented a fast (at ≈0.6 ns for SMAEA) and slow (at ≈125 ns for SMAQA) interaction between the polymer and amylin (initially separated by ≥1 nm) (Figure S9). As compared to SMAQA, SMAEA presented a higher number of intermolecular hydrogen bonds (H-bond), smaller secondary structure perturbation, lower root mean square deviation (RMSD), and lower root mean square fluctuation (RMSF) indicating a tight binding with amylin (Figures S10-S12). The RMSD plot presented a stable backbone dynamics for amylin in the presence of SMAQA, whereas SMAEA bound amylin showed short fluctuations during the last 150 ns (Figure S11a). Additionally, most of the amylin residues showed >4 Å RMSF in the SMAQA-amylin complex; whereas SMAEA-amylin complex selectively presented a comparatively small RMSF (average ∼ 2.7 Å) for residues 7 through 16 (Figure S11b) which is comprised of a α-helical structure. This indicates that the SMAEA binding to the central helix region could constrain amylin’s structural plasticity as compared to SMAQA interacting to the C-terminal domain. This is further confirmed by the secondary structure analysis of amylin as a function of time using Visual Molecular Dynamics program[36] (Figure S12) that exhibited an induction of α-helix (for residues 27-32) and several short transient β-sheets.

Atomistic interaction analysis (MD snapshot derived at every 100 ns) showed SMAEA initially binds to the N-terminal and C-terminal residues. SMAEA forms H-bonds with charged Lys1 residue following a structural rearrangement and the formation of a π-cation bond with His18 along with π-alkyl and hydrogen bonding to the central region spanning residues 13-18 (Figures S13 and S14). The π-cation interaction could be the major driving force to stabilize the α-helix, as has been shown to stabilize a protein structure by 0.4 kcal/mol.[37]

Atomistic interaction analysis (structure derived at 500 ns) showed SMAQA forming hydrophobic (Ile26, Val32 and Tyr37) and H-bonding (Asn31, Ser34, Asn35 and Thr36) interactions with amylin (Figure 5e, Table S1). SMAQA selectively binds to amylin’s C-terminal residues following an induction of a β-sheet structure in the N-terminus. Binding of SMAQA with one of the two β-sheets (residues 14-19 and 31-36) could be a possible mechanism of inhibition of amylin fibrillation.[38] Unlike SMAQA, SMAEA interacts with amylin (structure derived at 500 ns) at both N- and C-termini (Figure 5f, Table S1). Ala13 and Val17 showed hydrophobic packing, and the T-shaped π-π interaction was mediated by Phe23 (Figure 5f). The protonated His18 forms H-bond and electrostatic π-cation bond with SMAEA. Arg11 showed no interaction with SMAEA. However, a transient H-bond formation between SMAEA and Lys1 was observed during the first ∼350 ns (Figures S13 and S14). The electrostatic interaction by SMAEA could neutralize the positive charge on amylin in a manner similar to the effect of salt (Figure S6a) facilitating its quick aggregation as observed experimentally.[39] Since His-18 has been shown to coordinate with zinc metal ions, which influence amylin aggregation and fibril formation [40][41], it could be useful to investigate the interaction of SMAEA with human-IAPP in the presence of Zn^2+^ ions. We note that the structural maps generated from MD simulations on a microsecond regime may not be directly comparable with the reported experimental results considering the chemical and structural exchanges that happen in the millisecond-second regime. In addition, the results obtained from MD simulations could vary with the initial structure of IAPP monomers and applied force field in the MD calculation as documented elsewhere for intrinsically disordered proteins/peptides [42, 43]. Nevertheless, the molecular forces driving the peptide-polymer complex formation in different copolymers derived from MD simulations are useful to better understand the experimental observations reported in this study.

### Cytotoxicity of SMA copolymers and copolymer-induced amylin species

The toxicity of amylin species formed with SMA copolymers was measured both *in vitro* (HEK 293-T cells,) and *in vivo* (zebrafish embryo) experiments. Using MTT assay, we observed that SMA co-polymers are not toxic compared to cells treated with buffer only. In agreement with previous reports,[16, 44] samples formed by incubation of 10 µM of amylin monomers with or without continuous agitation at room temperature were cytotoxic. Samples of amylin co-incubated with 11 and 22 µg/mL SMA polymers were toxic compared to cells treated with 11 µg/mL SMA polymers (only) and buffer only (Figure 6a). Furthermore, cells treated with filtered (30 kDa MWCO) samples of amylin co-incubated with 22 µg/mL SMA co-polymers were not toxic compared to cells treated with buffer only (Figure 6a) suggesting that oligomers with a size larger than 30 kDa likely contribute to the toxicity. Although not statistically significant, and a higher concentration of amylin was used as compared to the physiological concentration, we find that amylin samples incubated without SMA polymers tend to be more toxic than amylin samples incubated with SMA polymers. Additionally, we would like to point out that amylin concentration in the blood plasma is different from that inside the cells that produce the hormone. In many instances, the concentrations of hormones inside the cells producing them can be orders of magnitude higher than the plasma levels. It should also be noted that the non-toxic nature of polymers (SMAEA or SMAQA) is encouraging for further development of these compounds into potential therapies.

**Figure 6.**
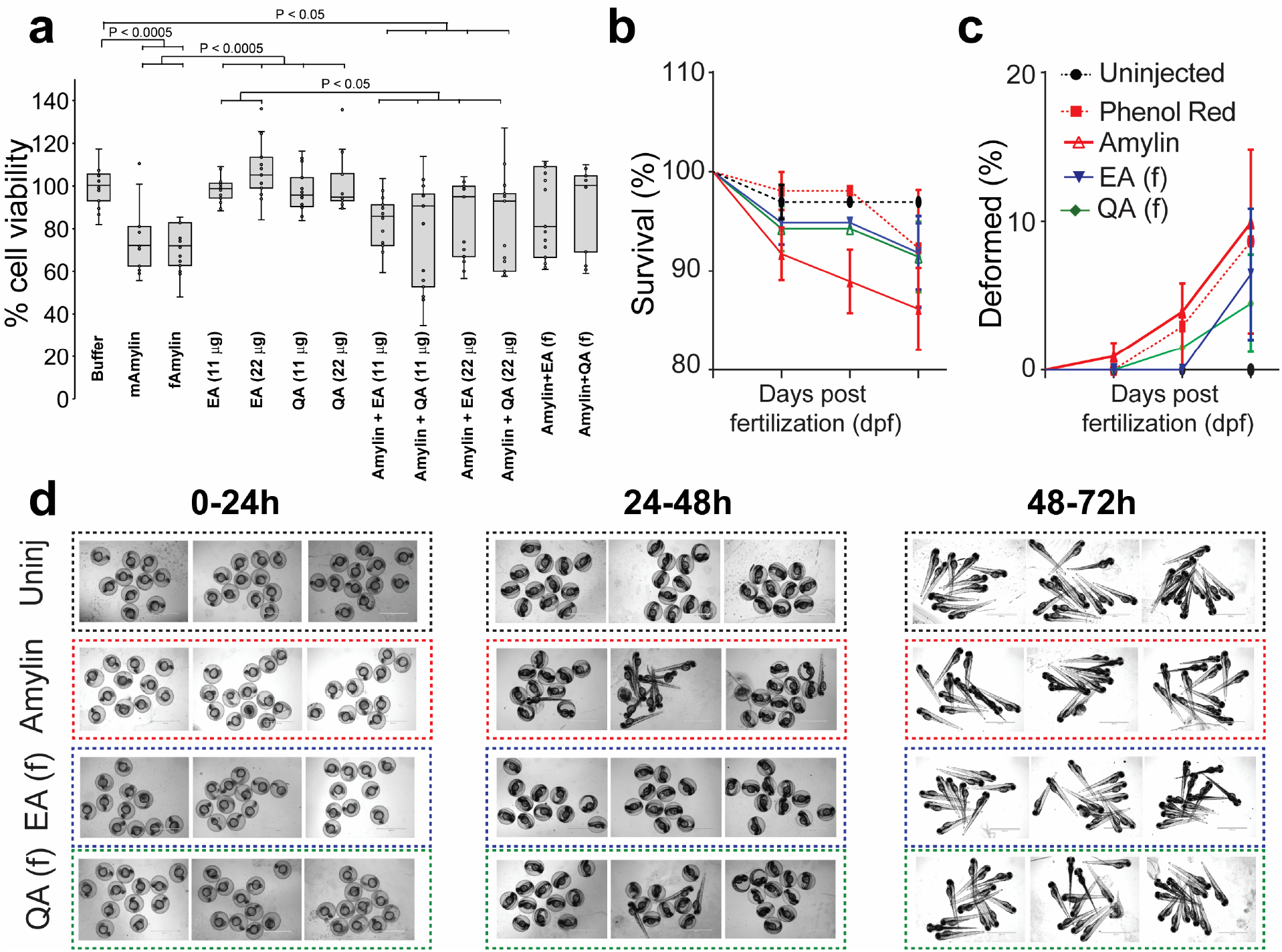
Modulation of amylin toxicity by SMA copolymers. (a) HEK-293T cell viability using MTT assay. Cells treated with SMA copolymer (EA or QA) incubated with/without 10µM of amylin at the indicated polymer concentrations. All samples were normalized using the MTT absorbance of cells incubated with 30mM NaAc buffer, pH 5.5. All quantitation results are shown as mean ± SEM for 3 technical (n) replicates and 5 independent (N) experiments; statistical analyses were done using Kruskal-Wallis test. The filtered amylin species obtained from the peptide-amylin mixture is denoted as EA (f) or QA (f). (b-d) % survival (b) and % deformity (c) of zebrafish embryo (N=3, n=15) treated with samples (10µM amylin; 22µg/mL polymers) indicated in color as a function of days-post-fertilization (dpf). (d) The statistical results are obtained from the images showing deformity in zebrafish embryo taken at different time-points. Scale= 2000 µm.

We also studied the effect of amylin samples formed with SMAEA and SMAQA zebrafish on the embryonic development. The successful delivery of the microinjected samples into the zebrafish embryo was first confirmed using phenol red. Zebrafish embryos injected with 11 or 22 µg/mL of SMAEA or SMAQA showed no significant mortality and deformity in the first 24 hours (Figure S15(a,b)). SMAEA (22 µg/mL) caused a decrease in survival rate (∼50 %) and deformity of embryos 48 hours post treatment indicating toxicity. In contrast, SMAQA (22 µg/mL) did not show a significant effect on zebrafish embryonic mortality and deformity 72 hours post treatment (Figure S15(a,b)). Both SMA copolymers were found to be not very toxic *in vivo* at least for 48 hours up to a concentration of 11 µg/mL which is relatively strong for zebrafish embryo. We tested this concentration considering the biophysical results used in this study; however *in vivo* concentration of amylin in diabetes patient is shown to be much lower (pg/mL).[45] 10 µM of amylin injected to zebrafish embryo showed a substantial mortality (equivalent to phenol red) and deformity 48- and 72-hour post-fertilization (Figures 6(b-d)). On the other hand, a relatively small percent of deformity was calculated for amylin incubated with SMAEA/SMAQA or for the filtered amylin species (Figure 6c) obtained from the polymer-peptide mixture (see methods). In particular, the filtered amylin species were shown to have the least toxic effect on the embryonic survival and deformity (Figure 6(b-d)).

## Conclusion

In conclusion, we have shown the distinct effects of two differently charged SMA-based copolymers on amylin aggregation as summarized in Table S2. The cationic SMAQA drives the formation of amylin globulomer, whereas the anionic SMAEA accelerates amyloid formation at physiological pH as well as under non-physiological ionic conditions. The reported biophysical results indicate that SMAEA-amylin interaction suppress the formation of amylin intermediates by quickly directing them to form β-sheet containing fibers. On the other hand, hydrogen bonding and hydrophobic interaction between SMAQA and amylin drives the formation of globular amylin intermediates. It is remarkable that this finding reflects the cationic amylin’s higher tendency to interact with anionic biomolecules such as anionic lipids resulting in the rapid formation of amyloids. On the contrary, hydrophobic driven interactions as observed for cationic amphiphilic, biomolecules (like SMAQA) restrict amylin’s fibrillation. Although a high peptide concentration was used in our study as compared to the physiological concentration, both SMA-based polymers showed a minimal cytotoxicity and their chemical properties could be modulated to control amylin toxicity. Although a combination of different biophysical techniques and toxicity measurements was used for a comprehensive investigation and to overcome the challenges in this study, the results obtained from different techniques may not be directly comparable due to the variation in the samples and experimental conditions used. On the other hand, the reported results can be used to develop further experimental strategies to solve atomic-resolution structures of amylin intermediates to better understand the correlation between structure and toxicity of amylin species. The generation of morphologically and pathologically distinct amylin species reported here by lipid solubilizing SMA copolymers[46–48] could be useful in strategically designing amyloid inhibitors that could potentially be useful to suppress IAPP induced cell toxicity in type-2 diabetes.

### Materials and MethodsSample preparation

All chemicals were purchased from Sigma Aldrich, with a purity greater than 95%. Synthesis and characterization of SMAEA and SMAQA polymers were carried out as reported elsewhere.[24, 25] Unless and otherwise mentioned, all samples were prepared using 30 mM sodium acetate (NaAc), pH 5.5. Synthetic human-amylin (KCNTATCATQRLANFLVHSSNNFGAILSSTNVGSNTY-NH_2_) was purchased from AnaSpec at >95% purity. The solid 1 mg amylin peptide was dissolved in 1 mL of 1,1,1,3,3,3- hexafluoroisopropanol (HFIP) and kept on ice for 30 minutes. Aliquots of 0.05 mg/mL amylin peptide dissolved in HFIP were next allowed to lyophilize for 48 hours. The dried powder was dissolved in NaAc buffer with a concentration ≤50µM and used immediately. Uniformly-^15^N isotope labelled human-amylin peptide was expressed in *E. coli* and purified following a previously reported protocol[34]. The expressed ^15^N amylin sample solutions were prepared in the same way as that of the synthetic amylin peptide.

### ThT fluorescence and critical micellar concentration (CMC) estimation

Thioflavin-T (ThT) based fluorescence assay was carried out to monitor the aggregation kinetics of the amylin peptide in the absence and presence of two different SMA-based polymers (anionic SMAEA and cationic SMAQA) following a previously published protocol.[12] Briefly, a 10 µM of freshly dissolved amylin peptide in NaAc buffer, pH 5.5, was added to a 96 well polystyrene plate with a total volume of 100 µL containing 10 µM of ThT dye and a variable concentration of polymer (5.5 to 44 µg/mL). ThT fluorescence measurement was also carried out under an identical condition as describe above for 10 µM of freshly dissolved amylin in the presence of 22 µg/mL of SMAEA dissolved in NaAc buffer containing a variable concentration (10 mM, 50 mM, 100 mM and 1 M) of sodium chloride. The aggregation kinetics was monitored at 25 °C for 72 hours under no shaking conditions for each sample in four replicates (N=4) using a Biotek Synergy-2 microplate reader with ThT excitation and emission wavelengths of 440 and 485 nm, respectively. Matured (or aged) amylin fiber was generated by incubating freshly dissolved monomers (100 µM) in NaAc buffer, pH 5.5 at room temperature under continuous agitation for ∼7 days. The relative fluorescence yield of 10 µM ThT in the presence of 11 or 22 µg/mL SMAEA was measured by titrating 1, 2 or 7 mol% (v/v) matured amylin fibers for 100 minutes using a 96-well plate reader at 25 °C. In addition to the ThT measurement of 10 µM amylin in the presence of copolymers under no-shaking conditions as described above, an experiment under medium shaking condition (5 µM amylin, 5.5 and 11 µg/mL copolymer) was also carried out at 25 °C.

The interference of SMAEA/SMAQA on ThT fluorescence yield was monitored using a FluoroMax 4® from HoribaScientific® in continuous mode at 25 °C. ThT was excited at 440 nm and emission spectra were collected from 450 to 700 nm (with a 5-nm bandwidth) with a 1 min delay using a 200-μL cuvette. Fluorescence spectra were recorded for 10 µM of ThT, 11 µg/mL SMAEA, 11 µg/mL SMAQA, 10 µM ThT mixed with 11 µg/mL SMAEA or SMAQA, 10 µM ThT mixed with freshly dissolved 5 µM amylin monomers or preformed matured amylin fibers. The matured amylin fibers were prepared from 100 µM amylin monomers under continuous agitation for ∼7 days. 11 µg/mL SMAEA or SMAQA mixed with 5 µM amylin monomers was incubated for ∼3 hours at room temperature. The sample mixture was next filtered using a 30-kDa Amicon® Ultra- centrifugal filter followed by washing (five times) using NaAc buffer, pH=5.5. The filtered samples were mixed with 10 µM ThT and then fluorescence spectra were collected. The binding affinity of filtered amylin globulomers with ThT dye was tested using a 96-well plate in triplicate under no-shaking condition at 25 °C as described above.

1 μM of pyrene solution was prepared from a stock of 0.1 mg/mL pyrene dissolved in 100% ethanol. SMAEA or SMAQA (10 mg/mL) was dissolved in NaAc buffer. A 200-μL cuvette containing different concentrations of copolymer and 1 μM of pyrene were used for fluorescence measurements. Pyrene was excited at 334 nm and the emission spectra were collected from 360 to 460 nm with a slit width of 2 nm using FluoroMax4® from Horiba Scientific® at 37 °C. The CMC value for each of the polymers was estimated from the ratio of pyrene’s fluorescence intensities of peaks I and III (I:III) by fitting the curve in Origin using a previously described method.[49]

### Circular Dichroism Spectroscopy and Fourier Transform Infrared spectroscopy

Far-UV circular dichroism (CD) spectrum of freshly dissolved 25 µM amylin monomers was recorded using a JASCO (J820) spectropolarimeter at 25 °C in the absence and presence of each of the copolymers. The CD spectra were recorded by subtracting the signals from buffer (for amylin) or buffer containing respective polymer (for amylin mixed with polymer samples). CD titration experiments were done by titrating 25 µM of amylin with a variable polymer concentration (11, 27.5 and 55 µg/mL) in NaAc buffer at pH 5.5. Next, the time-lapse CD measurements were carried out for the 25 µM of amylin in the absence or presence of 55 µg/mL of polymer incubated for 7 days at room temperature (day-2 refers to ∼24 hours of incubation) at buffer pH 5.5. CD spectra were also collected for samples containing 25 µM amylin dissolved in NaAc buffer, pH 8.5 in the absence or presence of 55 µg/mL SMAEA where amylin has a net charge of ≈0. In addition, CD samples were also prepared by dissolving 25 µM of amylin in NaAc buffer containing 0.25 M NaCl mixed with or without 55 µg/mL SMAEA. All CD spectra were averaged from a total of 16 scans and are presented as the mean residue ellipticity [Ɵ]. Secondary structure analyses were performed by the deconvolution of CD spectra using BeStsel.[29] Fourier Transform Infrared spectroscopy (FT-IR) spectra were recorded at 25 °C using a Thermos scientific ATR-FTIR instrument in transmission mode. Spectra were recorded within a range of 4000-400 cm^-1^ for the lyophilized sample powders. 25 µM of freshly dissolved amylin mixed with/without 55 µg/mL of SMAEA/SMAQA polymer was incubated for ∼12-hour at room temperature following lyophilization. Human-amylin fiber was prepared from 25 µM of freshly dissolved amylin monomers following continuous agitation at room temperature for ∼72 hours following lyophilization.

### Transmission electron microscopy

Transmission electron microscopy (TEM) imaging was obtained by incubating 25 µM of amylin mixed with 55 µg/mL of SMAEA or SMAQA for 24 hours at room temperature following a previously used protocol.[12] 20 μL of polymer mixed amylin sample was added to a collodion-coated copper grid and incubated for ∼4 minutes followed by three times rinsing with double deionized water. The grid was next stained with 4 μL of 2% (w/v) uranyl acetate, incubated for ∼2 minutes, rinsed three times with double deionized water followed by overnight drying under vacuum. TEM images were taken using a HITACHI H-7650 microscope (Hitachi, Tokyo, Japan).

### High-speed Atomic Force Microscopy

Real-time monitoring of amylin aggregation was performed using a HS-AFM instrument operated on tapping mode equipped with an Olympus BL- AC10DS-A2 cantilever of spring constant *k*=0.1 Nm^-1^ and frequency f=∼400 kHz in water. The instrument setup as described in a recently reported study.[12] Freshly dissolved amylin monomers (5 µM) dissolved in 30 mM NaAc buffer were incubated for ∼2-3 days at 37 °C under continuous agitation to form amyloid fibers. The amyloid seeds were generated by sonicating the fibers with a handheld sonicator and verified using AFM imaging (UR-21P, TOMY). The copolymer solutions of 11µg/mL were prepared in 30 mM NaAc buffer, pH 5.5. The fibril seeds were incubated on mica surface for ∼5 minutes, rinsed with buffer to remove the unbound seeds following immersion in 60 µL of buffer. The fibril seeds on mica surface were first confirmed using imaging and the 60 µL of buffer was replaced with 5 µM of freshly dissolved amylin monomers (fibril seeds:monomer=1:19 (v/v)) in the absence and presence of 11µg/mL of SMAEA or SMAQA following HS-AFM imaging. The HS-AFM videos were analyzed using ImageJ (NIH).

### NMR experiments

All proton NMR experiments were recorded on a 500 MHz Bruker NMR spectrometer equipped with a z-axis gradient triple resonance (TXO) probe at 25 °C. 2D NMR experiments were performed on a 800 MHz Bruker NMR spectrometer using a z-axis gradient cryogenic probe. ^1^H NMR spectra of 55 µg/mL of a copolymer, and 25 µM freshly dissolved amylin in the absence and presence of 55 µg/mL of a copolymer were acquired with 512 scans and a 2 s recycle delay. 2D ^15^N/^1^H SOFAST-HMQC NMR experiments[50] were carried out using 25 µM ^15^N amylin dissolved in 30 mM d3-NaAc, pH 5.5 containing 10% deuterated water in the absence and presence of 55 µg/mL of SMAQA or SMAEA at 25 °C. 2D NMR titration experiments were carried out with 32 scans and 256 t1 increments. Diffusion Ordered Spectroscopy (DOSY) and saturation transfer difference (STD) NMR experiments were carried out on a 500 MHz NMR spectrometer at 25 °C. DOSY and STD NMR spectra were recorded for 25 µM freshly dissolved amylin in NaAc buffer, pH 5.5 in the absence and presence of 55 µg/mL of SMAQA and SMAEA copolymer after ∼1 hour of incubation. DOSY experiments were obtained using stimulated-echo with bipolar gradient pulses for diffusion with gradient strength increment from 2 to 98%. 16 gradient strength increments, 36,000 time-domain data points in the t_2_ dimension, 3 s recycle delay, and 100 ms diffusion delay was used for DOSY measurements. DOSY spectra were analyzed using Bruker Topspin Dynamics center. STD NMR spectra were recorded with 256 number of scans, 4 s saturation time, an on-resonance excitation at -0.060 and 7.170 ppm (for amylin+SMAEA) and - 0.051, 3.078 and 7.256 ppm (for amylin+SMAQA), and off-resonance excitation at 40.0 ppm. ^1^H NMR spectra were also recorded for 25 µM amylin incubated with 55 µg/mL of SMAQA for ∼3 hr in d_3_NaAc buffer pH 5.5 following filtration (as described in section 2.2) and addition of 10% D_2_O. All NMR spectra were processed using TopSpin 3.5 (Bruker) and analyzed using Sparky.[51]

### Molecular dynamics simulations

The solution NMR structure (PDB: 5MGQ)[52] of human-amylin peptide was used for computational simulations. The amylin structure was initially processed using Gromacs *pdb2gmx* program with a total charge of +3.0 as appeared under physiological pH 5.5 condition. 3D PDB coordinates and topology parameter of SMAEA and SMAQA for the gromos54a7 force field were prepared by drawing the 2D structures of the copolymers followed by submission to Automated Topology Builder.[53] The SMA copolymer structure was energy minimized in GROMACS[54] prior to the preparation of peptide-polymer complex system for MD simulations. One molecule of amylin and one molecule of the polymer were placed ∼1 nm away from each other inside a cubic box of size 10×10×10 nm^3^ and solvated using SPC/E water model. The MD systems were neutralized using counter ions, energy minimized using steepest descent algorithm of tolerance of 1000 kJ mol^−1^ nm^−1^ followed by nvt and npt equilibration MD simulation as described previously[55] for 0.5 and 10 ns, respectively. A final production MD simulation was performed for 500 ns at 25 °C and the MD trajectories were analyzed using GROMACS local programs. All MD simulations were performed in GROMACS 5.0.7 using gromos54a7 force field running in parallel. MD snapshots were visualized using Discovery studio visualizer (Accelrys), Chimera, PyMOL and visual molecular dynamics (vmd).

### *In vitro* HEK-293 toxicity

HEK 293T/17 cells (ATCC® CRL-11268™) derived from human embryonic kidney were grown in in DMEM media with 2 mM L-glutamine supplemented with 10% fetal bovine serum at 37 °C in a humidified incubator with 5% CO_2_. Cells were kept between passage numbers 5-15. After splitting, 90 µL containing ∼35,000 cells were dispensed in a 96 flat-bottom and incubated for ∼24 hours prior to sample treatment. Samples were prepared by dissolving 100 µM amylin in 30 mM NaAc, pH 5.5 (referred as monomers) which were incubated without or with 11 or 22 µg/mL SMA copolymers at room temperature for ∼18 hours. To form aged fibrils, 100 µM of amylin monomers were incubated with shaking for ∼18 hours. Controls were cells treated with 30 mM NaAc buffer pH 5.5 (negative), 10% SDS containing 0.01M HCl (positive). All samples (10µL/well) were added in triplicate followed by incubation for ∼24 hours prior to cell viability assay. Statistical analysis were done from five independent experiments. Cell proliferation was measured by adding 10 µL/well of 12 mM MTT ((3-(4,5-Dimethylthiazol-2-yl)-2,5- Diphenyltetrazolium Bromide, CAS #298-93-1) stock. Following ∼1 hour incubation at 37 °C. 90µL of 10% SDS, 0.01M HCl solution was next added to each well. After ∼4 hours incubation, the absorbance was measured at 570 nm and 620 nm (background). Data was corrected by subtracting the background absorbance (620nm) following normalization with cells treated with SDS to 0% reduction, and cells treated with NaAc buffer to 100%. For statistical analysis, Kruskal-Wallis (KW) test was used.

### *In vivo* zebrafish toxicity

3% agarose molds were generated in advance using a mold (Adaptive Science Tools, TU-1) according to the manufacturer’s guidelines (http://site.adaptivesciencetools.com/Images/moldinstructions.pdf). Zebrafish adults were set up in breeding pairs overnight in breeder tanks. At 8 a.m., dividers were pulled and fish were monitored for egg production. 1-hour post-fertilization embryos were injected with 2nL of each construct over a 3-hour period. Microinjections were done for all groups that include Sham, phenol red, 10µM amylin mixed with or without 22 µg/mL SMA copolymers, 22 µg/mL SMA copolymers, and filtered amylin species obtained from polymer-peptide (22 µg/mL polymer and 10 µM amylin) mixture (see method section 2.2) using a micromanipulator and glass needles (Sutter Instrument, Novato, CA) following established protocol.[56] The needles were replaced between each group to prevent cross-contamination of experimental conditions. Microinjections for the SMA copolymers (22 µg/mL) were done using two beakers (N=2) each containing 15 number of embryos (n=15). Other groups such as amylin, amylin+SMA copolymers, filtered amylin species and phenol red were carried out using N=3 and n=15. Zebrafish embryos were next washed with sterile-filtered Embryo Rearing Media and transferred to a 26 °C incubator. Deformity and mortalities were recorded every 24hrs for 72hrs, followed by the removal of dead embryos, 90% media change, and photographs of each group by EVOS light microscope at 4X magnification.

## Acknowledgements

This study was supported by NIH (AG048934 to A.R.) and the cell toxicity assay was supported by NIH (NS096785-10 to MII) and the computations at the Osaka University through the Institute for protein research “International Collaborative Research” (ICR-18-02) program.

