## Supporting information for "Conformational Tuning of Amylin by Charged Styrene-maleic-acid Copolymers"

**Table of Content**

1. Supporting Tables (Table 1 and 2)
2. Supporting figures (Figures S1-S15)
3. Supporting movies (Movies SV1-SV6)
4. References

**Table S1.** Interaction analysis between SMA copolymers and amylin derived from 500 ns all-atom MD simulation snapshot.

| SMA-amylin complex | Bond type |
|---|---|
| SMAEA - THR9 | Hydrogen Bond |
| SMAEA - ALA13 | π-Alkyl |
| SMAEA - ASN14 | Hydrogen Bond |
| SMAEA - VAL17 | π-Alkyl |
| SMAEA - HIS18 | Electrostatic (π-Cation) |
| SMAEA - PHE23 | π-π T-shaped |
| SMAEA - GLY24 | Hydrogen Bond |
| SMAEA - SER29 | Hydrogen Bond |
| SMAEA – THR30 | Hydrogen Bond |
| SMAEA – ASN31 | Hydrogen Bond |
| SMAQA - ILE26 | π-Alkyl |
| SMAQA - ASN31 | Hydrogen Bond |
| SMAQA - VAL32 | π-Alkyl |
| SMAQA - SER34 | Hydrogen Bond |
| SMAQA - ASN35 | Hydrogen Bond |
| SMAQA – THR36 | Hydrogen Bond |
| SMAQA - TYR37 | π-Sigma |

| Amylin behavior | in the presence of SMAEA | in the presence of SMAQA | Method(s) |
|---|---|---|---|
| Aggregation kinetics | Accelerate fibrillation at high-concentration | Delay fibrillation at high-concentration | Fluorescence spectroscopy |
| ThT interference | Yes | No | Fluorescence assay |
| Secondary structure | Induce β-sheet structure | Induce α-helix structure | CD spectroscopy and FT-IR |
| Morphology | Amyloid fiber | Globular oligomer | HS-AFM and TEM |
| Fibril growth in presence of seeds | Yes | No | HS-AFM |
| SMA copolymer binding region | Both N- and C-termini | C-terminus | Atomistic MD simulation |
| Interaction (intermolecular H-bond formation) | Fast (~0.6 ns) | Slow (~125 ns) | Atomistic MD simulation |
| Reduction in NMR signal intensity | Most of the residues (amide) are not detectable | Most of the residues (amide) are detectable | SOFAST-HMQC and $^{1}$H NMR |
| Cell toxicity | Nontoxic | Nontoxic | In vitro assay |
| ---- | Nontoxic | Nontoxic | In vivo |

**Table S2.** A comparative summary of the observed differences in amylin behavior in the presence of SMAEA or SMAQA.

### 2. Supporting figures

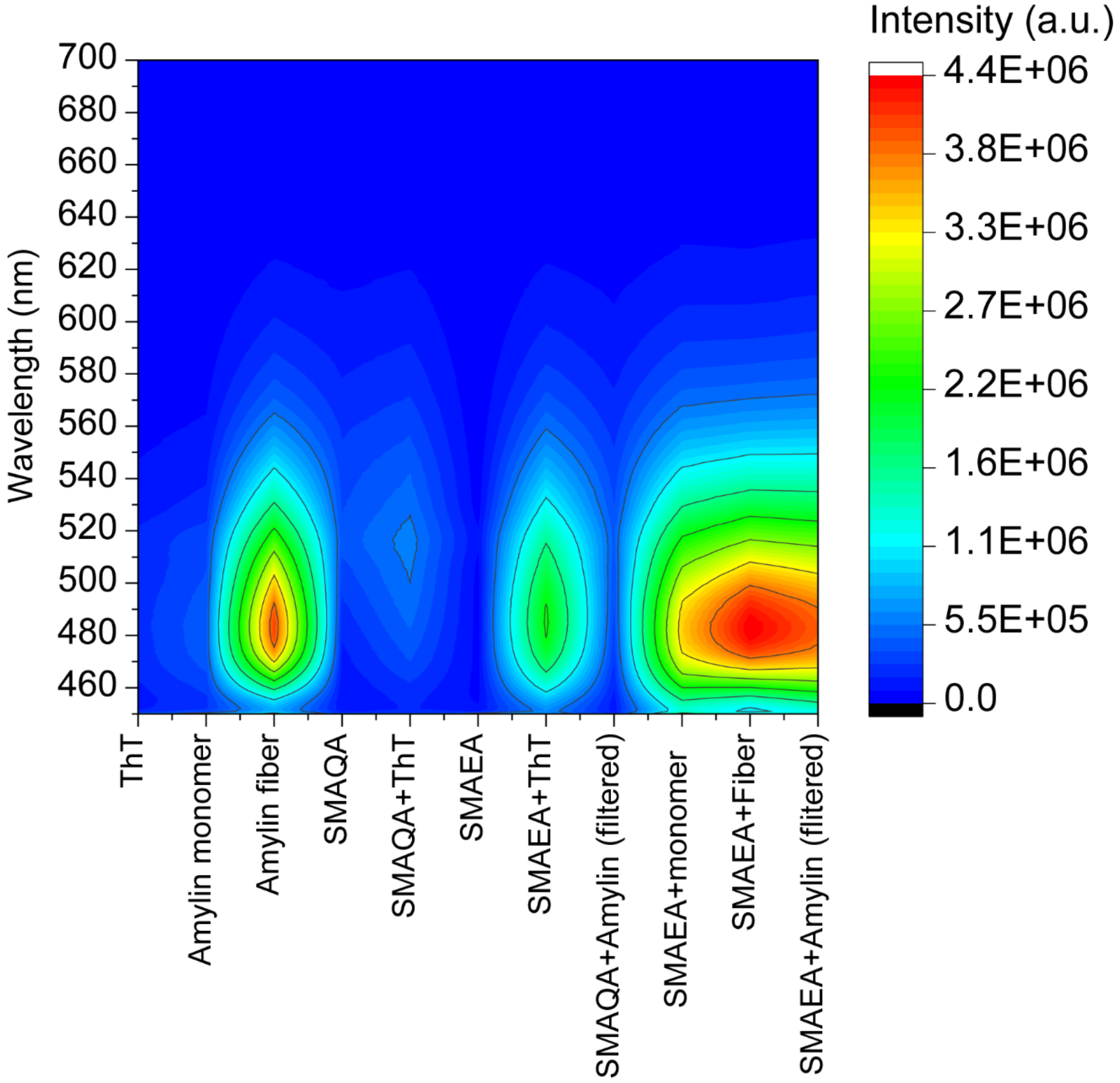

**Figure S1.** 2D plot of the fluorescence emission of 10 μM thioflavin-T (ThT) dye mixed with and without different polymer/peptide mixture as indicated. The SMAEA/SMAQA and amylin concentrations used for this study were 11 μg/mL and 5 μM, respectively. Aged amylin fibers were generated using freshly dissolved amylin monomers (see methods). The polymer+amylin mixed sample was filtered using a 30-kDa Amicon ultra centrifuge filter (see methods), and fluorescence measured for these samples are referred as 'filtered'. ThT fluorescence was measured at an excitation wavelength of 440 nm and the emission spectrum was recorded from 450-700 nm at 25 °C. All measurements were carried out using a 200 μL quartz cuvette in 30 mM NaAc buffer, pH 5.5. The concentration of amylin fiber (100 μM stock) used for fluorescence was 5% v/v.

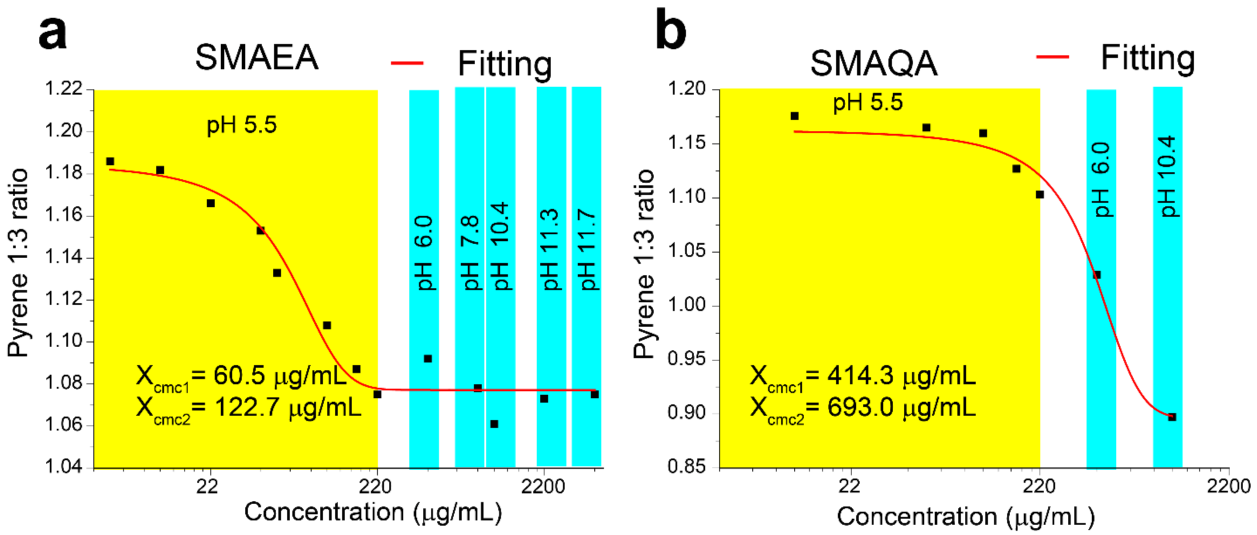

**Figure S2.** Critical micellar concentration (cmc) of SMAEA (a) and SMAQA (b) determined using pyrene assay. The polymer was dissolved in NaAc buffer, pH 5.5; and the change in pH during titration is highlighted. The cmc plot was generated from the ratio of the intensities of I and III peaks measured from the fluorescence spectra of pyrene (pyrene 1:3 ratio) and fitted using the Origin program. $X_{cmc1}$ and $X_{cmc2}$ denotes the CMC1 and CMC2 as described previously.[1]

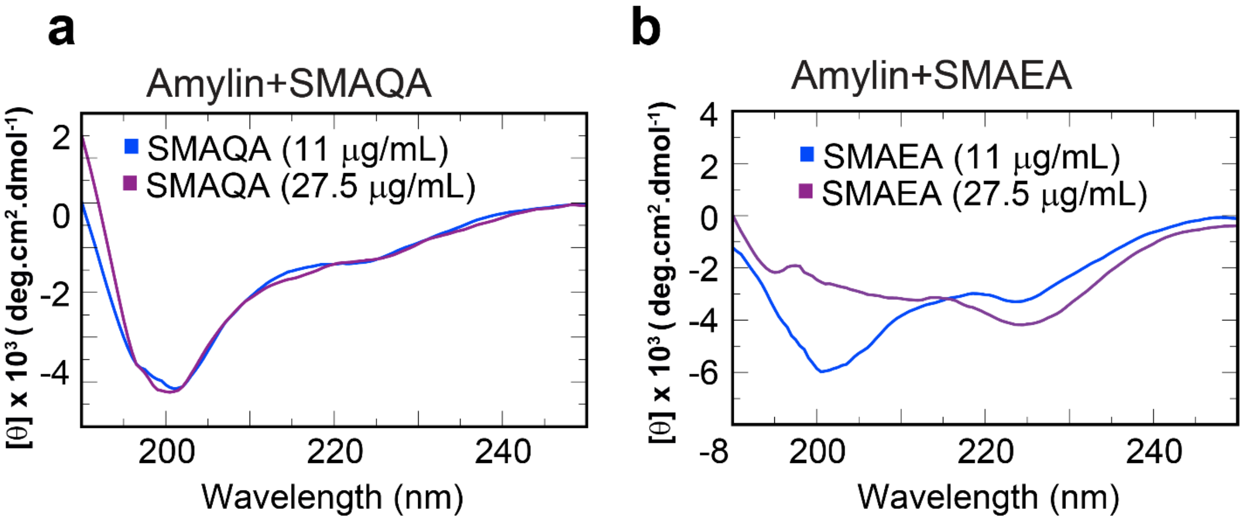

**Figure S3.** Far-UV CD spectra of 25 μM of human amylin mixed with SMAQA (a) and SMAEA (b) at the indicated polymer concentration recorded at 25℃.

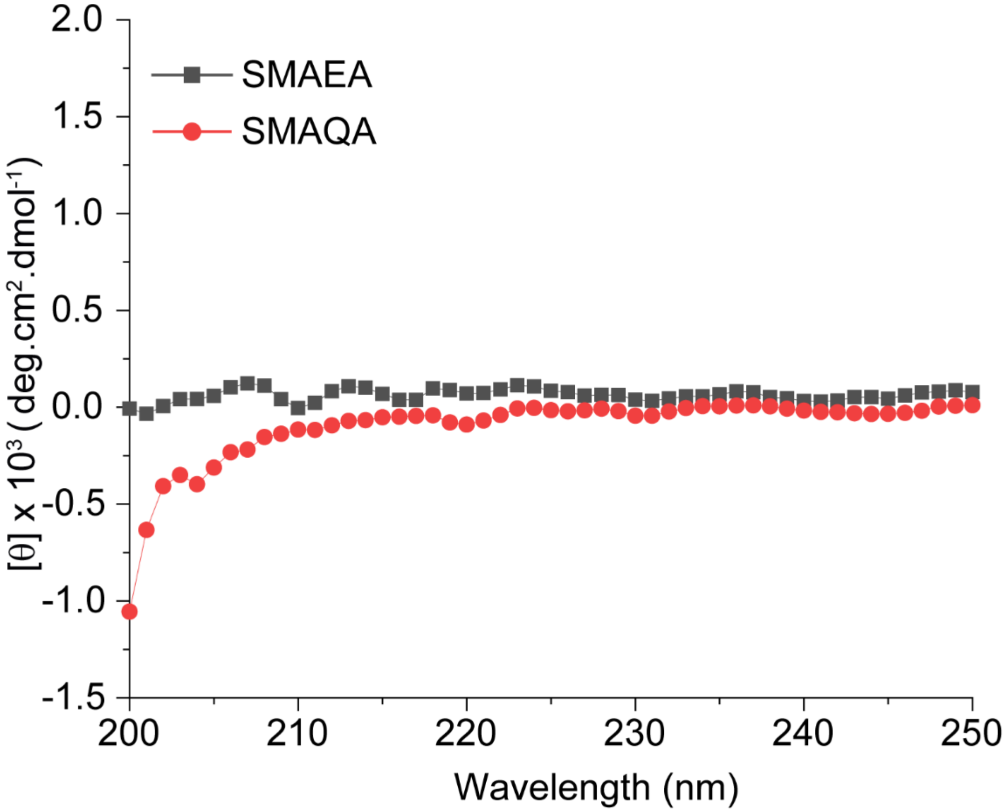

**Figure S4.** Far-UV CD spectra of 55 µg/mL of SMAEA (black) and SMAQA (red) dissolved in 30 mM NaAc buffer, pH 5.5 at the indicated colors.

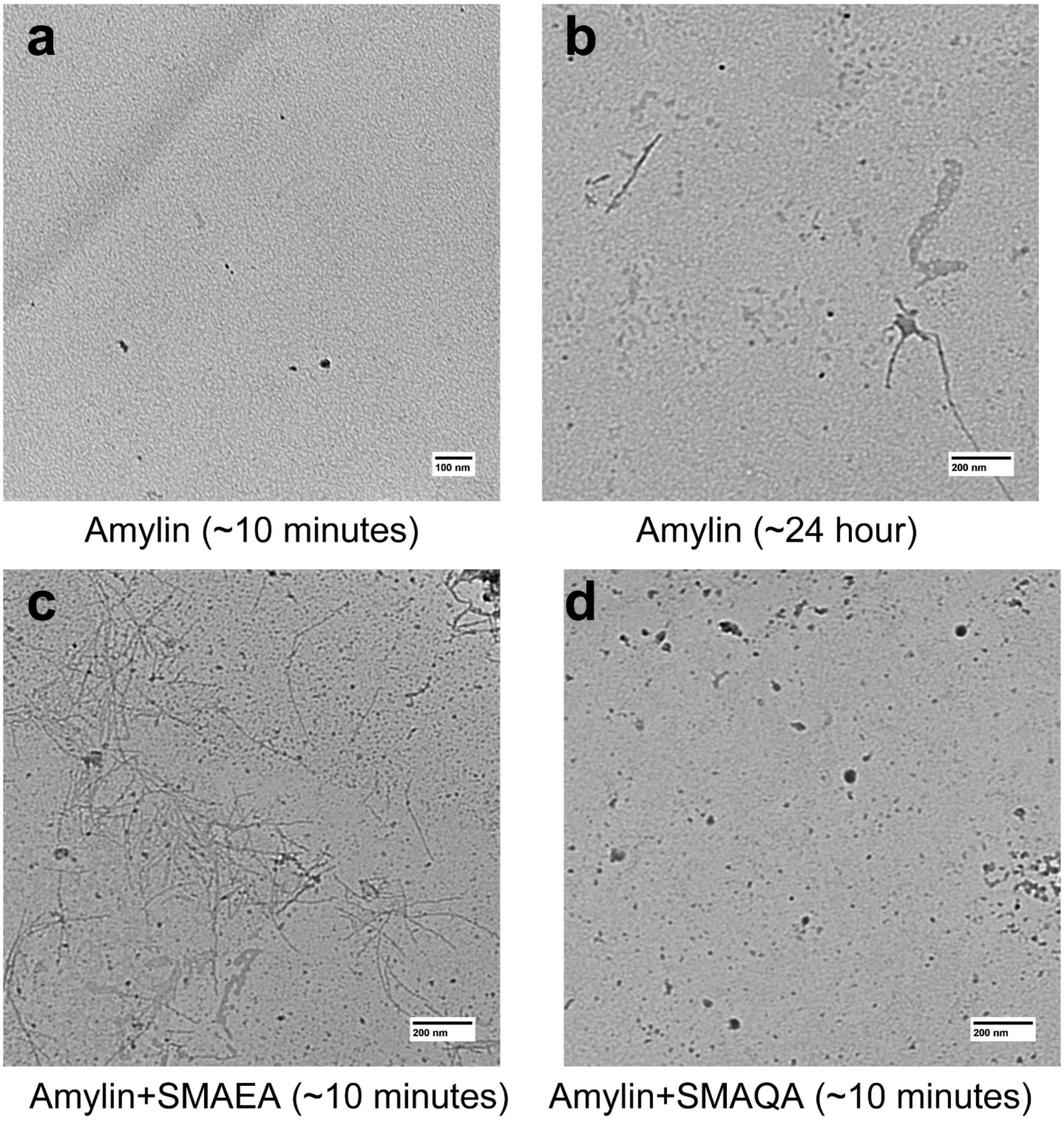

**Figure S5.** Negative staining TEM images of 25 μM human amylin incubated for the indicated times without (a and b) and with 55 μg/mL of SMAEA (c) or 55 μg/mL SMAQA (d) at 25℃. The scale bar is 100 nm for (a) and 200 nm for (b-d).

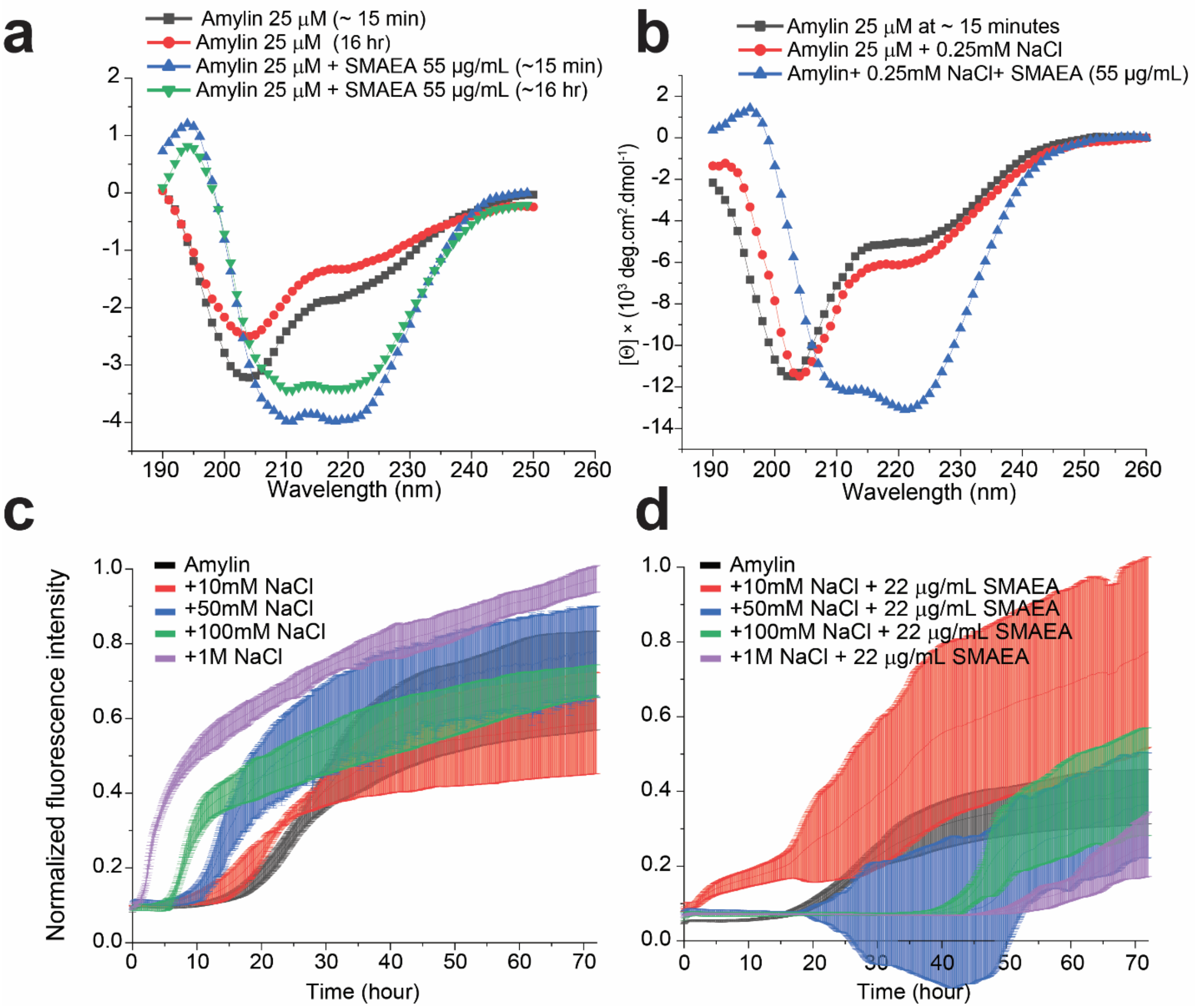

**Figure S6.** (a) Time-lapse Far-UV CD spectra of 25 µM of human amylin mixed with 55 µg/mL of SMAEA dissolved in NaAc buffer, pH 8.5 recorded at 25℃. (b) Far-UV CD spectra of 25 µM of human amylin mixed with 55 µg/mL of SMAEA dissolved in NaAc buffer, pH 5.5 containing 0.25 mM NaCl recorded at 25℃. (c-d) Normalized ThT fluorescence (N=4) of 10 µM amylin in the presence of a variable concentration of NaCl (c), and in the presence of 22 μg/mL SMAEA and variable amount of NaCl (d) as indicated in color.

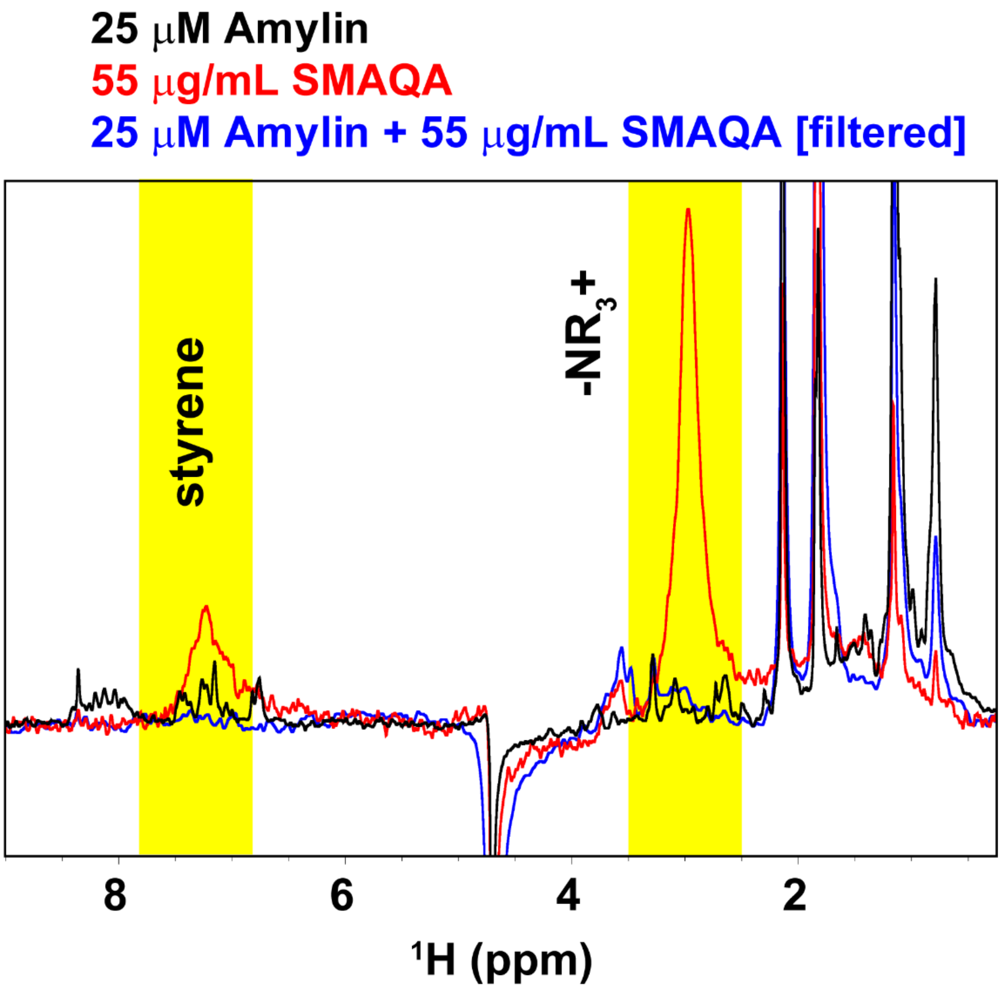

**Figure S7.** 25 μM of human amylin dissolved in $d_3$-NaAc buffer was mixed with 55 μg/mL of SMAQA and incubated overnight at room temperature following filtration using a 30 kDa Amicon Ultra centrifugal filter. 10% $D_2O$ was added to the filtered sample following proton NMR measurement (blue spectrum) at 25 °C. NMR measurements were carried out on a 500 MHz Bruker NMR spectrometer. For comparison, $^1$H NMR spectra of 25 μM of human amylin monomer (black) or 55 μg/mL of SMAQA (red) dissolved in $d_3$-NaAc buffer were measured. The yellow-highlighted highlighted regions show the unique SMAQA peaks that are absent in the filtered sample indicating the absence of SMAQA.

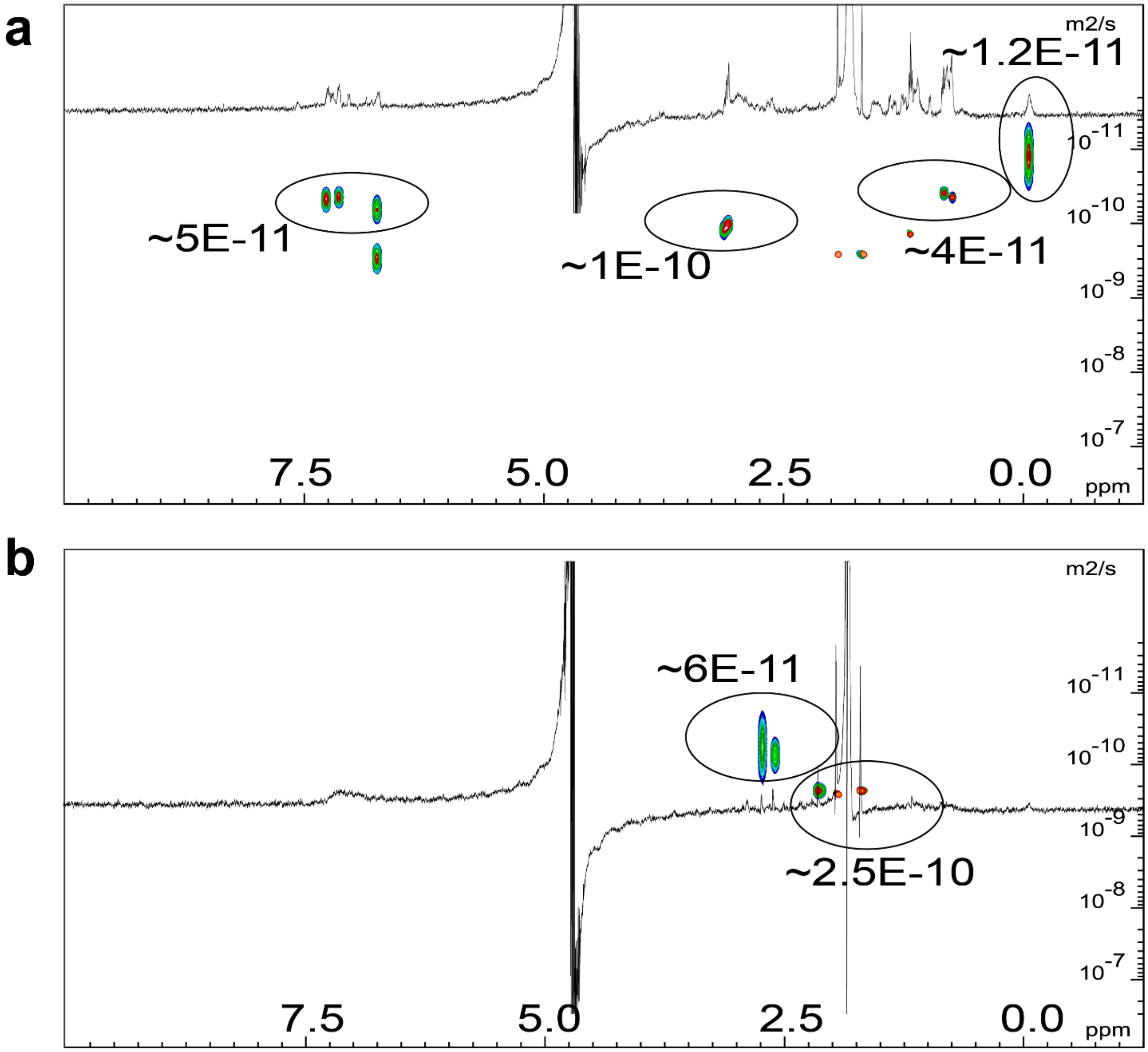

**Figure S8.** 2D DOSY spectrum of 25 μM amylin in the presence of 55 μg/mL SMAQA (a) or SMAEA (b) incubated for ≈ 1 hour at room temperature. The NMR spectra were recorded on a 500 MHz Bruker NMR spectrometer at 25 °C. The diffusion constants (in $m^2/s$) and corresponding molecular species are indicated for a few selected peaks and the 1D spectrum is shown inside each 2D plot.

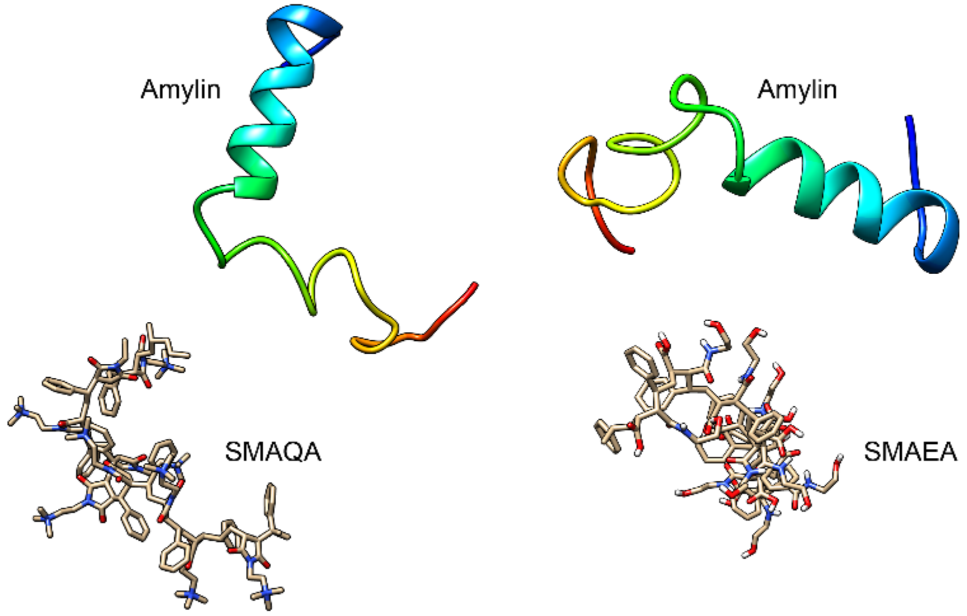

**Figure S9.** Snapshots showing the molecular dynamics setup and initial conformations of human amylin (PDB: 5MGQ) and SMA copolymers in a cubic box of size 10x10x10 $nm^3$. The peptide and polymer molecules were initially separated by a distance ≥ 1 nm.

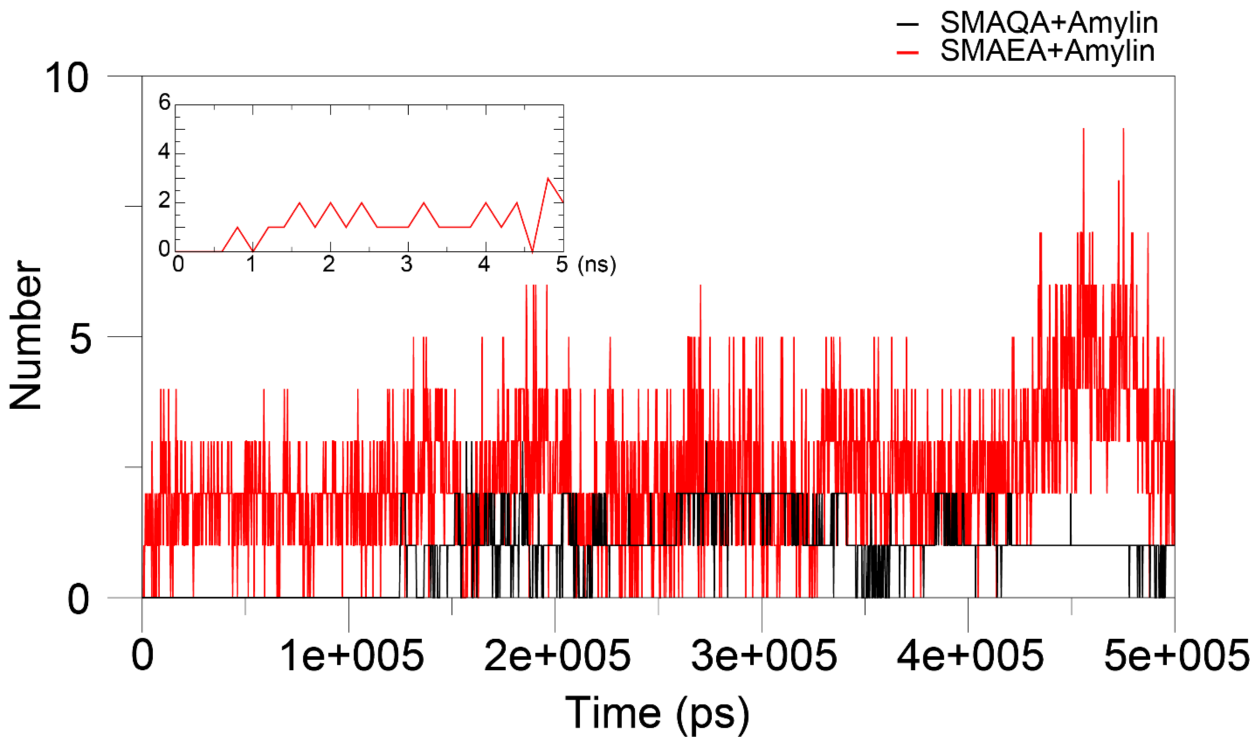

**Figure S10.** Graph shows the number of hydrogen bonds between human amylin and SMAEA/SMAQA copolymer derived from 500 ns MD simulation as a function of time. The inset shows the formation of hydrogen bonds between SMAEA and human amylin at ≈600 ns.

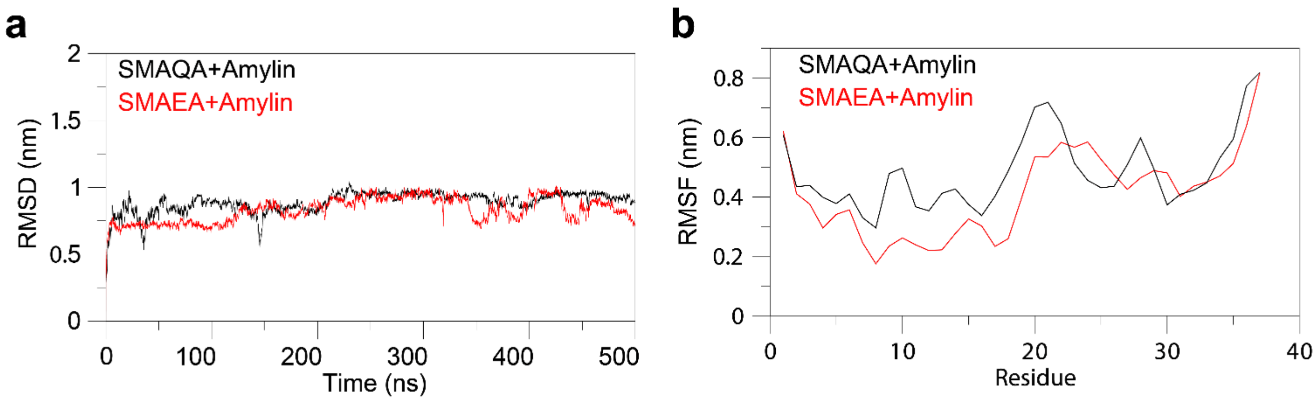

**Figure S11.** (a) Root mean square deviation (RMSD) of human amylin backbone atoms in the amylin-SMA copolymer complex as a function of time derived from 500 ns MD simulations. (b) Root mean square fluctuations (RMSF) of human amylin residues derived from the amylin-SMA copolymer complex as a function of amino acids.

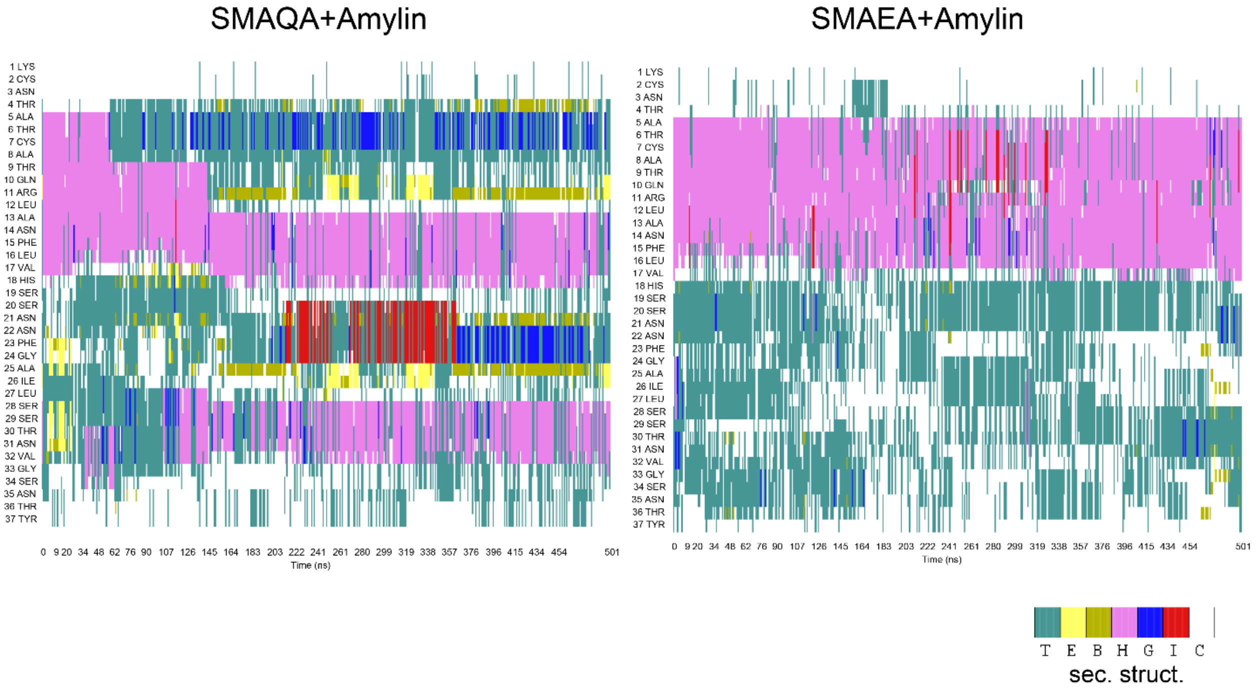

**Figure S12.** Secondary structure evolution map of human amylin derived from 500 ns MD simulations in the amylin-SMA copolymer complexes. The secondary structure units are shown at the bottom, where T: Turns, E: Extended β-strand, B: Isolated bridge, H: α-helix, G:3-10 helix, I: Pi-helix and C: coil.

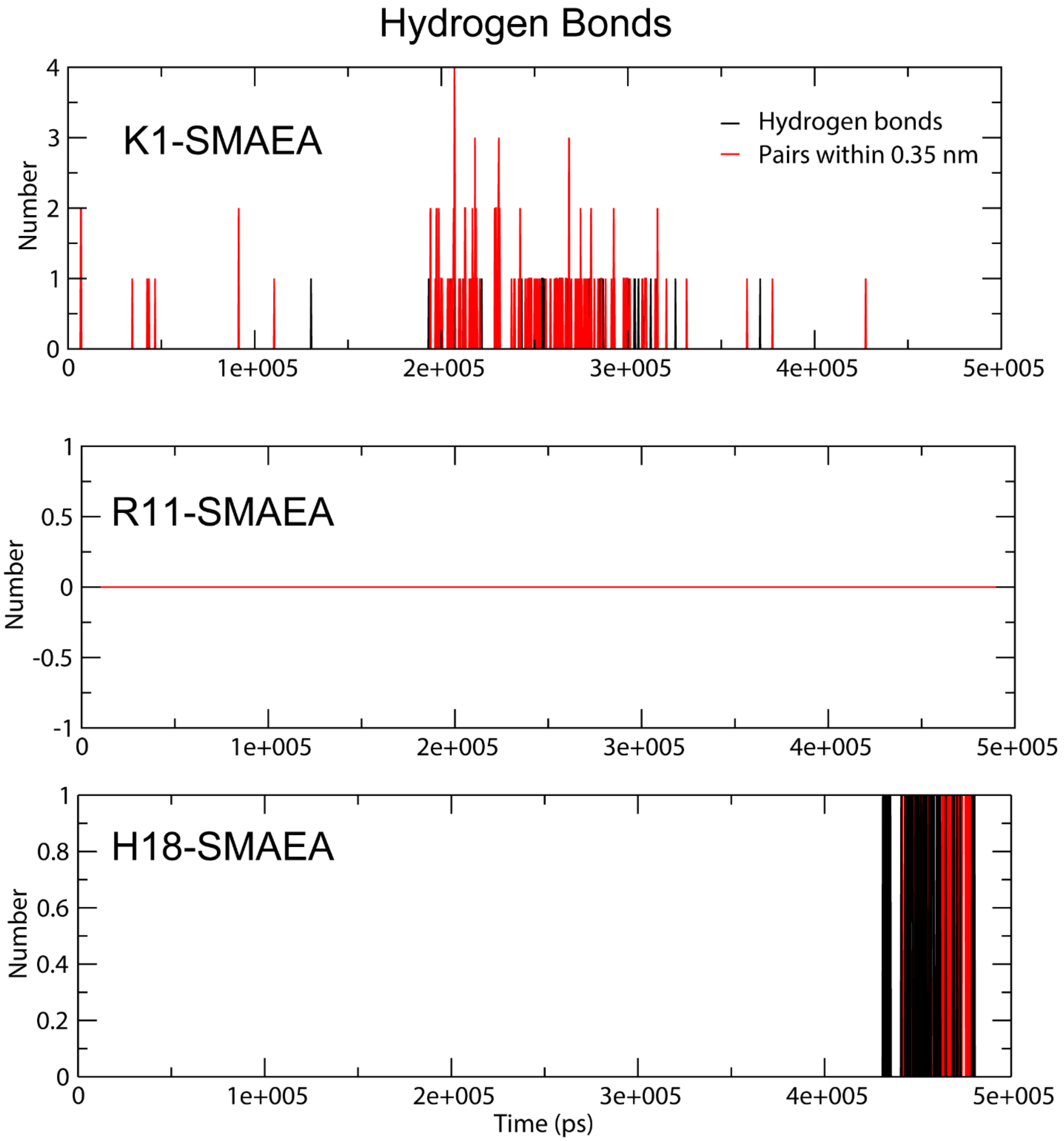

**Figure S13.** Graph shows the number of hydrogen bonds between the positively charged human amylin residues (Lys1, Arg11 and His18) and a SMA copolymer derived from 500 ns MD simulation as a function of time.

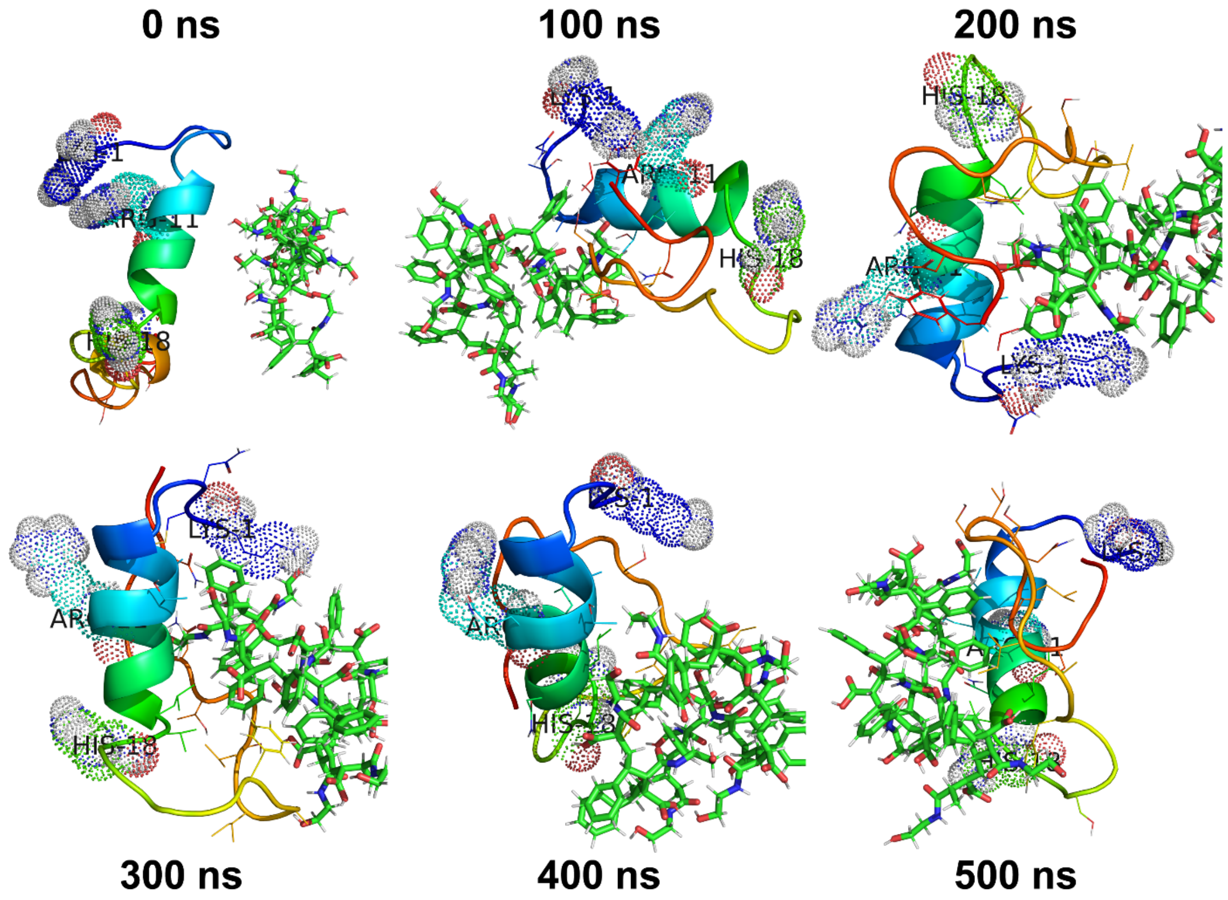

**Figure S14.** MD snapshots of the SMAEA-amylin complex retrieved at every 100 ns from the 500 ns MD simulation. The amylin peptide is shown in *cartoon*, SMAEA in *sticks*, and charged residues Lys1, Arg11 and His18 in *dots* in PyMOL.

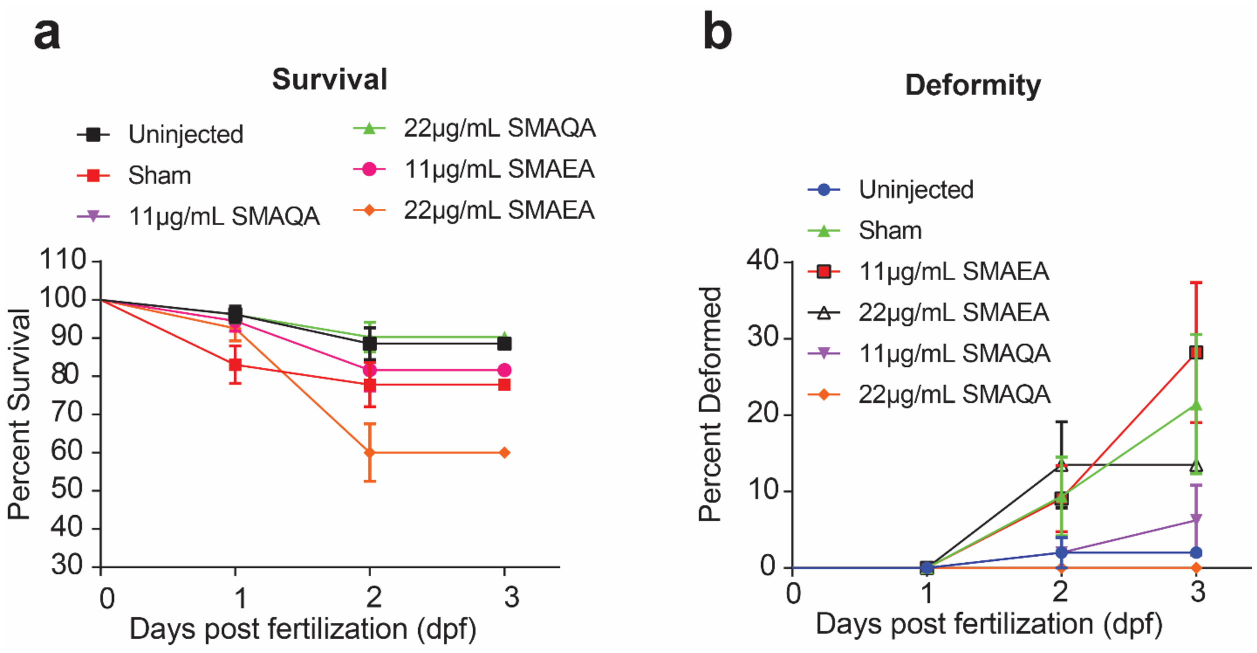

**Figure S15.** (a) Effect of SMAEA and SMAQA on zebrafish embryo survival at the indicated colors relative to the uninjected embryos. The percent deformity in zebrafish embryos treated with SMA copolymers (b) and 10 µM amylin in the presence or absence of SMA copolymers as indicated at 24, 48, and 72 hours post-fertilization (dpf). The standard errors shown were obtained from N=2 (two beakers) and n=15 (number of zebrafish embryos).

### 3. Supporting movies

**3.1 Movie SV1.** Real-time monitoring of 5 µM human amylin fibrillation growth in the presence of amylin seeds (fibril seeds:monomer=1:19).

**3.2 Movie SV2.** Real-time monitoring of 5 µM human amylin fibrillation growth in the presence of 5.5 µg/mL of SMAQA and amylin seeds (fibril seeds:monomer=1:19).

**3.3 Movie SV3.** HS-AFM showing the formation of *de novo* human amylin globumers in the presence of SMAQA. The size and growth of the *de novo* globumers are plotted by calculating the grey value where 1 pixel=2 nm as shown in Figure 1b and d.

**3.4 Movie SV4.** HS-AFM showing the growth of an isolated fibril of human amylin in the presence of amylin seeds (fibril seeds:monomer=1:19).

**3.5 Movies SV5.** Real-time monitoring of 5 µM human amylin fibrillation growth in the presence of 5.5 µg/mL of SMAEA and amylin seeds (fibril seeds:monomer=1:19).

**3.6 Movie SV6.** HS-AFM showing the growth of a *de novo* human amylin globumer (marked inside a circle) to form fibers in the presence of 5.5 µg/mL of SMAEA and amylin seeds (fibril seeds:monomer=1:19).
